# More than black and white: complex relationships involving serine proteases regulate the Toll pathway and the melanization response in *Drosophila*

**DOI:** 10.1101/383257

**Authors:** J.P. Dudzic, M.A. Hanson, I. latsenko, S. Kondo, B. Lemaitre

## Abstract

The melanization response is a rapid and important defense mechanism in arthropods. Melanin is produced around wound sites and invading microorganisms by phenoloxidases (POs), which need to be activated by the sequential activation of an extracellular serine protease (SP) cascade. *Drosophila melanogaster* has been a useful genetic model for dissecting insect immune signaling, but understanding these proteolytic cascades has been complicated by the large number of SP genes, possibly with redundant function. Taking advantage of recently-generated null and compound mutants, we re-investigated the role of SPs involved in the melanization response in *D. melanogaster* and discovered phenotypes previously concealed in single mutant analysis. We found that two of them, Hayan and Sp7, can activate the melanization response in two different manners: Hayan is required in the local blackening of wound sites, while Sp7 regulates an alternate melanization reaction responsible for the clearance of septic infections with *Staphylococcus aureus*. We present evidence that both Sp7 and Hayan regulate the Toll NF-κB pathway. Sp7 is regulated by canonical Toll signaling downstream of PGRP-SA, ModSP, and Grass, leading to control of septic infections via a Sp7-dependent melanization response. Additionally, we found that Hayan and the Toll-regulating SP Psh are the result of a recent gene duplication. Using genetic manipulations, we reveal the hidden role for Hayan, alongside Psh, in propagating Toll signaling downstream of pattern recognition receptors. Thus, we describe the existence of two pathways leading to the melanization response and reveal previously unknown dynamics in the activation of the Toll pathway.

## Introduction

Sequential activation of extracellular serine protease (SP) cascades regulates important aspects of insect innate immune reactions, notably activation of the Toll pathway and the melanization response. These proteolytic cascades have a functional core, consisting of several SPs that undergo zymogen activation upon cleavage by an upstream protease. This sequential cleavage shapes the immune response by providing a link between recognition and the induction of effectors (Lemaitre and Hoffmann, 2007). In *Drosophila*, the Toll pathway mediates resistance to Gram-positive bacteria and fungi by regulating a subset of antimicrobial peptides in the fat body. Unlike mammalian Toll-like receptors that function as pattern recognition receptors, the *Drosophila* Toll receptor does not interact directly with microbial determinants and is instead activated by a cleaved form of the secreted molecule Spätzle (Spz) (Lemaitre et al., 1996; Weber et al., 2003). The immune-regulated CLIP-domain SP Spätzle processing enzyme (SPE) has been identified as the terminal SP that cleaves Spz (Jang et al., 2006). Genetic analysis supports the existence of two complex cascades that link microbial recognition to activation of SPE: the pattern recognition receptor (PRR) and Persephone (Psh) pathways. In the PRR pathway, PRRs involved in the sensing of Gram-positive bacteria (GNBP1 or PGRP-SA) or fungi (GNBP3) bind to their respective microbial ligands to activate an upstream SP, ModSP, which then leads to the activation of the SP Grass, leading to the maturation of SPE (Buchon et al., 2009; Chamy et al., 2008; Gobert, 2003; Gottar et al., 2006). In the Psh pathway, infectious agents are detected directly through the cleavage of the protease bait region of the SP Psh by microbial proteases, leading to Spz cleavage by SPE (Issa et al., 2018).

Melanization is one of the most spectacular immune reactions in insects. It is an arthropod-specific immune response resulting in the rapid deposition of the black pigment melanin at wound or infection sites (Cerenius et al., 2008; González-Santoyo and Córdoba-Aguilar, 2012; Tang, 2009). This process relies on enzymes called prophenoloxidases (PPOs), which catalyze the oxidation of phenols resulting in the polymerization of melanin. The microbicidal mechanism by which melanization contributes to the killing of bacteria, fungi or parasitoid wasp larvae remains elusive, although reports indicate a role for reactive oxygen species and other metabolic intermediates of the melanin synthesis pathway (reviewed in Nappi et al., 2009). In *D. melanogaster*, three PPOs have been identified. PPO1 and PPO2 are produced by crystal cells and contribute to hemolymph melanization, while the role of the lamellocyte-derived PPO3 is confined to encapsulation (Binggeli et al., 2014; Dudzic et al., 2015; Nam et al., 2008). Thus far, three SPs have been implicated in activating PPOs in the hemolymph: MP1, Sp7 and Hayan (Castillejo-López and Häcker, 2005; Nam et al., 2012; Tang et al., 2006). While a null mutation in *Hayan* totally abolishes melanization in adults, a null *Sp7* mutation results in only a slight reduction (Dudzic et al., 2015). The positions of these SPs in the melanization cascade has not been fully established, although Hayan and Sp7 have been shown to cleave PPO1 *in vivo* and *in vitro* (An et al., 2013; Nam et al., 2012). It is still unclear whether PPO1 and PPO2 are differentially activated by the same or by distinct SP cascades.

In many insects, the Toll and melanization pathways are activated by the same SPs, diverting only at the terminal steps (Kan et al., 2008; Park et al., 2006; Volz et al., 2006). However, a common opinion is that the SPs regulating the Toll pathway in *Drosophila* upon infection are independent of the melanization cascade and vice versa. Nevertheless, the melanization and Toll pathways interact, as the Toll pathway regulates the expression of many genes encoding SPs and serpins involved in the melanization pathway (De Gregorio, 2002; De Gregorio et al., 2002; Ligoxygakis, 2002). Moreover, there has been disparate evidence that PRRs upstream of Toll, such as PGRP-SA and GNBP1, can impact Toll-independent responses, notably melanization (Matskevich et al., 2010).

In this article, we introduce a new mode of systemic infection using a low dose of *Staphylococcus aureus* (*S. aureus*) where the survival of flies relies entirely on the melanization response, but not on expression of antimicrobial peptides or phagocytosis. Using this sensitive assay, we show that resistance to *S. aureus* correlates with the melanization response, but surprisingly not the deposition of melanin itself. We also reveal specific roles for the SPs Sp7 and Hayan in the melanization pathway: Hayan is specifically tied to the blackening of the cuticle, while the Sp7-dependent melanization response is mediated by Toll PRR signaling, which diverges at Grass to activate either SPE or Sp7. Meanwhile, a small deficiency removing both *Hayan* and *psh* reveals an unexpected role for these two SPs in the canonical Toll pathway. We provide evidence that *Hayan* and *psh* arose from a recent gene duplication, and that these genes currently have both overlapping and distinct functions in the melanization and Toll pathways. Globally, we describe the existence of two distinct pathways leading to melanization, and reveal a previously unappreciated role for both Hayan and Psh in Toll signaling.

## Results

### Melanization is important to survive *Staphylococcus aureus* infection

While reviewing survival data from Binggeli et al. (2014), we confirmed that survival of *D. melanogaster* adult flies against *S. aureus* is strongly dependent on a functional melanization response. Flies lacking *PPO1* rapidly succumb to infection with a low dose of *S. aureus* (OD 0.5, 1-100 CFU/fly), whereas the lack of *PPO2* alone is not critical. A synergistic effect can be observed in flies lacking both *PPO* genes simultaneously; in this case no fly survives the infection (Fig. 1A). Analysis of the bacterial load of *PPO1^Δ^,2^Δ^* flies reveals that melanization-deficient flies already have a higher bacterial burden 6 hours (h) after infection compared to their wild-type (*w^1118^*) counterparts (Fig. 1B). Their inability to control *S. aureus* growth is even more prominent 24 h after infection. Following injection of GFP-producing *S. aureus, PPO1^Δ^,2^Δ^* flies do not melanize the wound (Fig. 1C arrows) and exhibit a local GFP signal after 18 h, indicating *S. aureus* growth at the wound area (Fig. 1C). After 24 h, most *PPO1^Δ^,2^Δ^* flies exhibit a strong, systemic GFP signal indicating that they were unable to suppress *S. aureus* growth and spread (Fig. 1C). This contrasts with wild-type flies, which deposit melanin at the wound area and control bacterial growth over 24 h (Fig. 1D). The importance of the melanization response to resist *S. aureus* infections is further illustrated by the observation that *PPO1^Δ^,2^Δ^* flies show no defect in Toll or IMD pathway activation after *S. aureus* infection. Compared to wild-type flies, they even show a ~7x higher expression of the Toll-activity readout *Drosomycin* (*Drs*) as well as ~8x higher expression of the IMD-activity readout *Diptericin* (*Dpt*) (Fig. S1A,B), likely due to unconstrained bacterial growth. To rule out the involvement of hemocytes in the resistance to *S. aureus*, we produced hemoless flies using a plasmatocyte-specific gal4 (*hml^Δ^-Gal4*) to express the pro-apoptotic gene *bax* (Defaye et al., 2009), and additionally examined *eater*-deficient flies who exhibit a defect in *S. aureus* phagocytosis (Bretscher et al., 2015). In both cases, hemoless and *eater* deficient flies survived like the wild-type (Fig. S1C). Our results contrast with previously published data showing that hemocytes are critical to survive *S. aureus* infections via phagocytosis (Defaye et al., 2009; Garg and Wu, 2014). We attribute this discrepancy to using a low dose of *S. aureus* that is controlled specifically by the melanization response but not phagocytes.

**Fig. 1:**
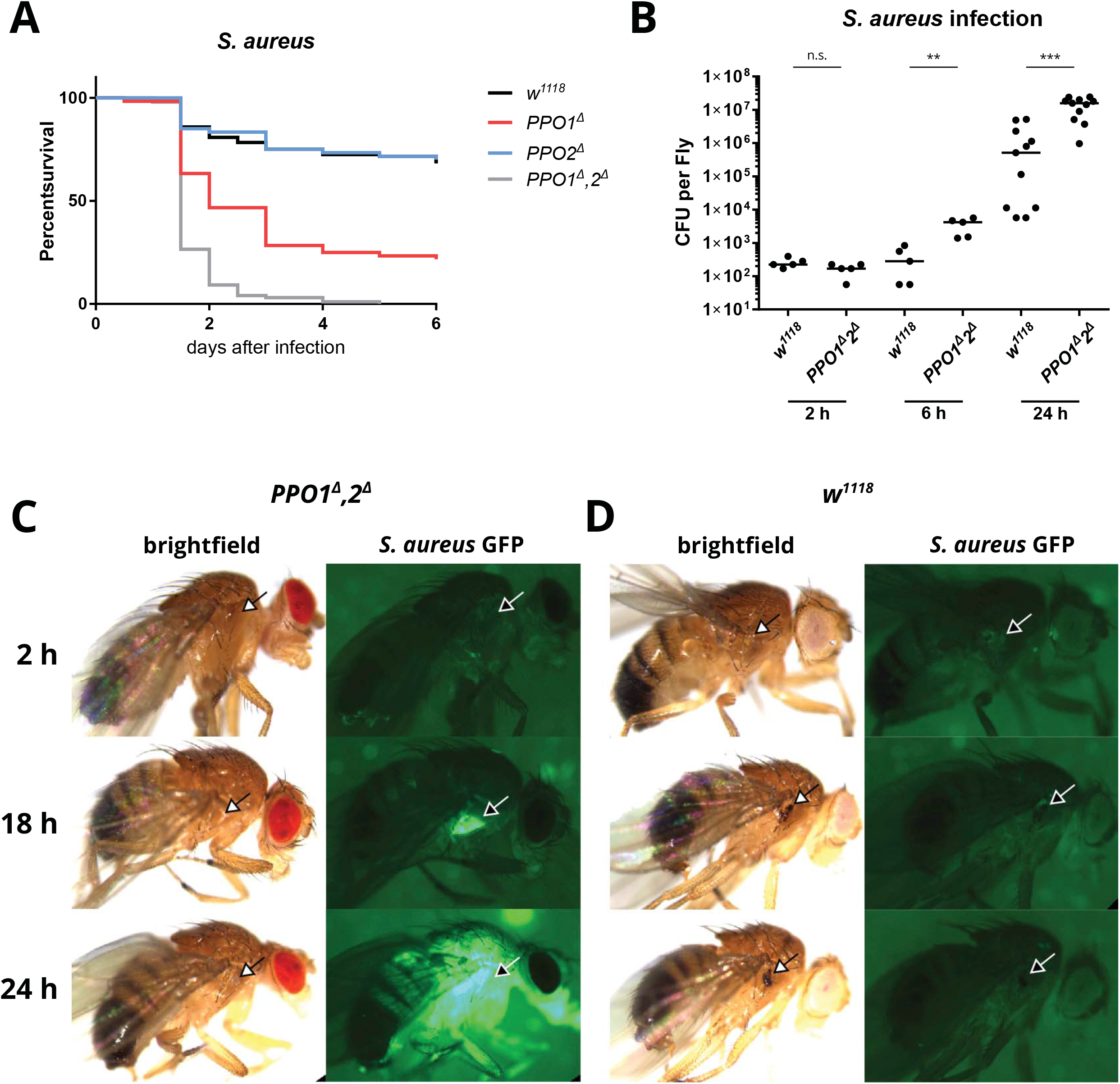
PPOs contribute to the resistance of *Staphylococcus aureus*. **A:** Survival rates of flies following septic injury with Gram-positive *S. aureus*. Flies lacking *PPO1* (P < 0.0001) but not *PPO2* (n.s.) alone are less resistant than wild-type flies (w^1118^). Flies lacking two PPO genes (*PPO1^Δ^,2^Δ^*) (P < 0.0001) show less resistance than flies lacking only *PPO1*. **B:** Persistence of *S. aureus* in *w^1118^* or *PPO1^Δ^,2^Δ^* flies at 2, 6 or 24 hours post-infection. Increased *S. aureus* counts are found in *PPO1^Δ^,2^Δ^* flies after 6 and 24 hours. The number of colony forming units (CFU) per fly is shown on a logarithmic scale. Data were analyzed by Mann-Whitney test and significance is indicated in the graph. **C,D:** Growth of GFP expressing *S. aureus* in *PPO1^Δ^,2^Δ^* (C) or *w^1118^* (D) flies after 2, 18 and 24 hour*S. Wild-type* flies melanize the wound area (arrows) while *PPO1^Δ^,2^Δ^* do not. No GFP signal is observed in *w^1118^* flies while *PPO1^Δ^,2^Δ^* exhibit a local GFP signal after 18 hours, and systemic GFP signal after 24 hours. Exemplary micrographs are shown.

Cytotoxic by-products of the melanization reaction include reactive oxygen species (ROS) (Nappi et al., 2009). We investigated the role of ROS during *S. aureus* infections by measuring H2O2 levels in whole fly lysates using a fluorimetric approach. We did not see a change in H2O2 levels over a time-span of 6 hours following *S. aureus* infection in wild-type flies. Interestingly, *PPO1^Δ^,2^Δ^* flies show a non-significant but consistent reduction in H_2_O_2_ levels compared to wild-type flies (Fig. S1D).

Taken together, these results demonstrate that the melanization response is critical to resist a low dose of *S. aureus* infection consistent with a previous study (Binggeli et al., 2014). Furthermore, our *S. aureus* infection model provides a sensitive assay to characterize the role of melanization in resistance to infection.

### Sp7 but not Hayan nor MP1 are required to resist *S. aureus* infection

Using this low dose *S. aureus* infection model, we analyzed the role of three SPs (MP1, Sp7 and Hayan) that were previously described to be involved in melanization. For this, we used a newly generated CRISPR null mutant for *MP1* (*MP1^SK6^*, this study), and null mutants for *Sp7* and *Hayan* (*Sp7^SK6^, Hayan^SK3^*, Dudzic et al., 2015). Contradicting a previous report that used an *in vivo RNAi* approach (Tang et al., 2006), we did not observe any overt defect in Toll and PPO activation in *MP1* deficient flies, whose immune phenotype characterization is further described in Fig. S2. We therefore focused our attention on the respective roles of Hayan and Sp7. Surprisingly, only *Sp7* mutants phenocopy the susceptibility of *PPO1^Δ^,2^Δ^* flies against *S. aureus*, while *Hayan* deficient flies behave like wild-type (Fig. 2A). By injecting GFP-producing *S. aureus* in flies lacking these serine proteases, we observed that *Sp7^SK6^* flies fail to restrict *S. aureus* growth first locally after 18 h, then systemically after 24 h (Fig. 2C, arrows). In contrast, *Hayan^SK3^* flies show no increase in GFP signal over time, indicating that *S. aureus* proliferation is controlled as in the wild type (Fig. 2D, arrows).

**Fig. 2:**
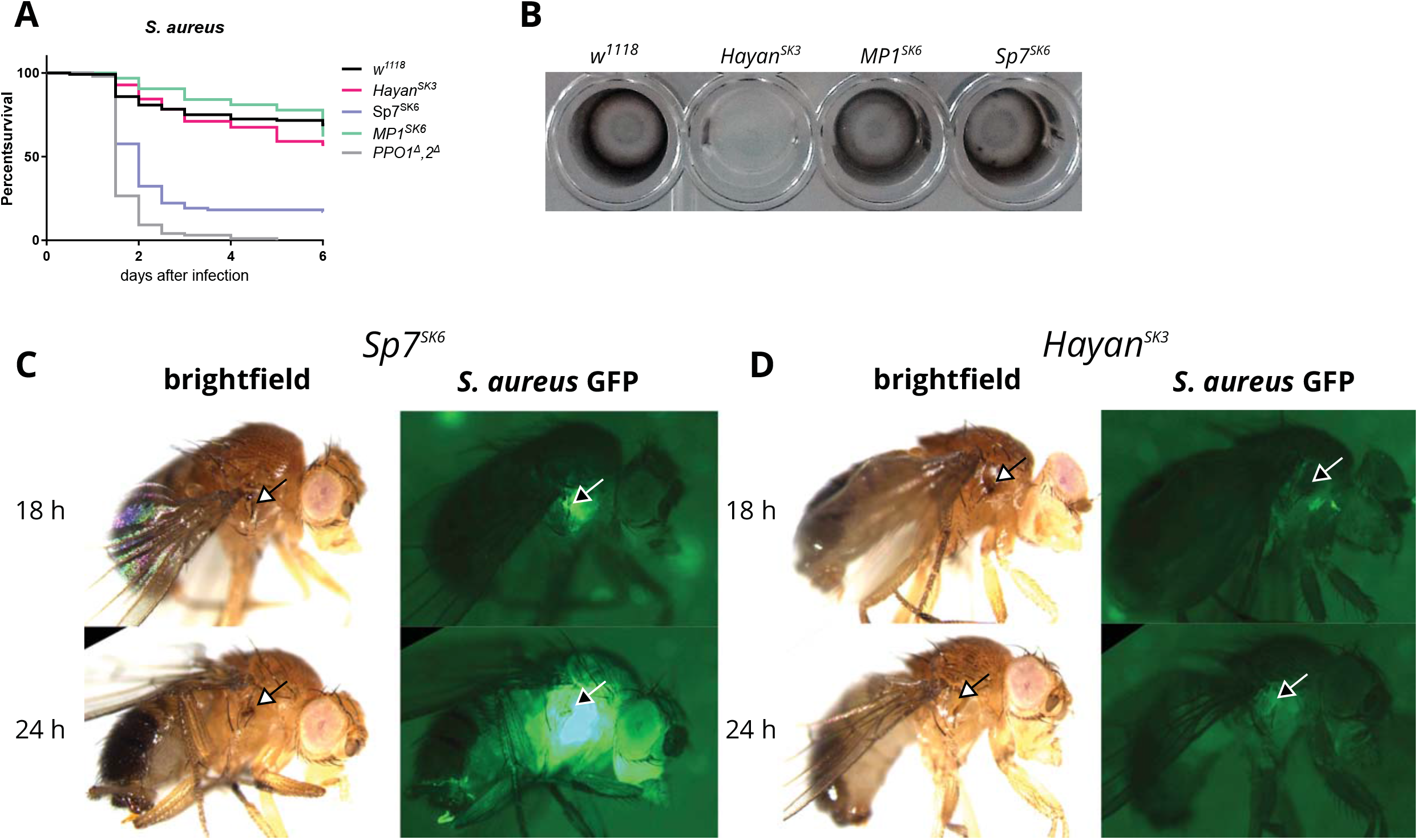
*Sp7* but not *Hayan* or *MP1* is critical to resist *S. aureus* infections. **A:** Survival rates of flies following septic injury with Gram-positive *S. aureus*. Flies lacking *Sp7* (P < 0.0001) but not *Hayan* (P = 0.0075) or *MP1* (P = 0.3156) are less resistant than wild-type flies (w^1118^). *PPO1^Δ^,2^Δ^* flies were used as a positive control. P-values are from log-rank tests compared to wild-type flies. **B:** Capability of L3 larval hemolymph to produce black melanin after incubation at room temperature. Hemolymph from *w^1118^, MP1^SK6^* and *Sp7^SK6^* flies shows melanization while *Hayan^SK3^* hemolymph fails to melanize. **C, D:** Growth of GFP producing *S. aureus* in *Sp7^SK6^* (B) or *Hayan^SK3^* (C) flies after 18 and 24 hour*S. Sp7^SK6^* flies show a local GFP signal at the wound area (arrows) while *Hayan^SK3^* flies show a strong reduction. In *Sp7* mutants, this moves into a systemic GFP signal after 24 hours. Exemplary micrographs are shown.

These results were surprising since *Hayan*, but not *Sp7* mutants have reduced melanization at the wound site (hereon referred to as the “blackening” reaction) after clean injury (CI) (Fig. S2A) and *S. aureus* (Fig. 2C,D) (Dudzic et al., 2015; Nam et al., 2012) We then investigated the role of Sp7 and Hayan in the melanization of the hemolymph in both adults and larvae. Extracting hemolymph with subsequent incubation at room temperature leads to blackening, due to PO-dependent melanin production. Surprisingly, hemolymph from both adults and larvae shows that *Hayan^SK3^* mutants fail to produce melanin, while hemolymph of *Sp7^SK6^* mutants turns black over time (Fig. 2B for larval hemolymph, data not shown for adults). Together, these results disconnect the blackening of the hemolymph from a melanization reaction-dependent clearance of *S. aureus*, suggesting that intermediate metabolites in the melanization reaction contribute to microbial control.

Collectively, our results show that Sp7, but not Hayan, controls *S. aureus* and that the underlying resistance mechanism does not involve melanin deposition, a terminal step in the melanization cascade. This suggests that by-products of the melanization reaction, such as ROS or other metabolic intermediates, might be the active molecules controlling bacterial growth. We also reveal the complexity of the PPO cascade in *Drosophila*, pointing to the existence of multiple branches involving either Sp7 or Hayan. To disentangle the deposition of melanin from the melanization reaction as a whole, we will use ‘blackening reaction’ to refer to melanin deposition and ‘melanization response’ to refer to PPO-derived activities as a whole, which includes the blackening reaction.

### Sp7 and Hayan regulate PPOs differently

Melanization is a reaction that can be triggered by clean injury or by the presence of microbial products. To further explore the relationship between the blackening reaction and the two SPs, we compared the blackening reaction at the wound site upon clean injury or septic injury with the avirulent Gram-positive bacterium *Micrococcus luteus*. For this, we categorized the blackening reaction of the cuticle into three levels: strong, weak and none. An example for each category is given in Fig. S3. For wild-type *w^1118^* flies, after clean injury 91.8% blacken strongly and 8.2% blacken weakly. This ratio shifts slightly for *Sp7* mutants (66.6% strong, 30.6% weak, 2.8% none, p = 0.008, Fig. 3A grey columns), while *Hayan* mutants were almost deficient for the blackening reaction (0% strong, 38.9% weak, 61.1% none, p < 0.0001). Interestingly, some cuticle blackening is recovered in *Hayan* mutants upon septic injury with *M. luteus* (16.2% strong, 64.6% weak, 19.2% none, p < 0.0001, Fig. 3A red columns). Thus, Hayan is not solely responsible for the blackening reaction, as the presence of bacteria can induce cuticle blackening in a Hayan-independent manner.

**Fig. 3:**
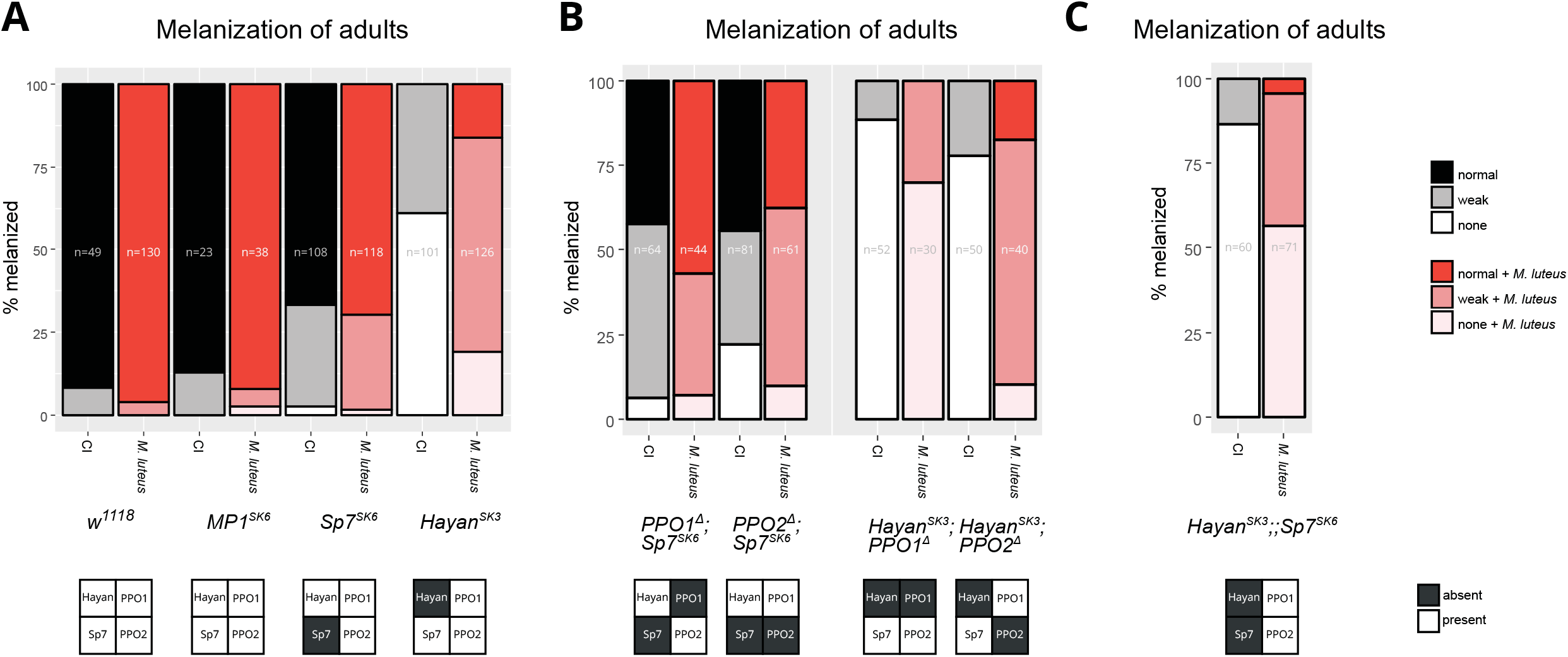
Sp7 and Hayan regulate PPOs differently. Flies were either injured with a clean needle (black bars) or with a needle previously dipped in *M. luteus* solution (red bars). Melanization was assessed in three categories: strong, weak or none (see Fig. S2). **A:** *MP1^SK6^* flies do not exhibit any defect in melanization compared to *w^1118^* flies (p = 0.8219). *Sp7^SK6^* flies show a reduction in strong melanization although almost all flies melanize at least weakly (p = 0.0083). *Hayan^SK3^* flies show a strong defect in melanization after clean injury (p < 0.0001), which can partially be rescued by wounding with *M. luteus* (p < 0.0001). **B:** *Sp7* mutants still melanize the wound area to a certain extent with a simultaneous mutation in either *PPO1* or *PPO2*. In contrast, the partial melanization of *Hayan^SK3^* flies after infection with *M. luteus* relies on the presence of *PPO1*. **C:** *Hayan^SK3^;;Sp7^SK6^* mutants lose most, but not all melanization. Percentages of total flies (n) are displayed. Sample size (n) is indicated in each respective bar. Data analyzed with Pearson’s Chi-squared.

In *Drosophila*, two PPOs, PPO1 and PPO2, produce the bulk of hemolymph PO activity as no blackening is observed in *PPO1, PPO2* deficient mutants. However, it is unclear whether these PPOs are activated by the same SP or are differentially regulated. To further understand how the SPs Hayan and Sp7 relate to PPO1 and PPO2, we generated double mutants and subsequently analyzed their melanization capabilities upon clean injury and *M. luteus* infection. Previous reports have shown that mutations in *PPO1*, and to a lesser extent *PPO2*, reduce melanization upon both clean and septic injury (Binggeli et al., 2014). Here, *PPO1^Δ^,Sp7^SK6^* and *PPO2^Δ^,Sp7^SK6^* retain a blackening reaction, only slightly reduced compared to *PPO1^Δ^ or PPO2^Δ^* alone (data not shown). This indicates that Sp7 is not essential for the blackening reaction, and that Hayan (or another SP) can activate either PPO1 or PPO2 in the absence of Sp7 (Fig. 3B). In contrast, *Hayan^SK3^;PPO1^Δ^* double mutants fail to blacken, even upon septic injury with *M. luteus*, which is not the case for *Hayan^SK3^;PPO2^Δ^* (Fig. 3B). As Hayan is not responsible for the additional blackening we see upon septic injury (Fig. 3A red), this indicates that another SP acts on PPO1, possibly Sp7, after exposure to Gram-positive bacteria. Finally, we observed that flies, double mutant for both *Hayan* and *Sp7* have strongly reduced but still detectable levels of the blackening reaction (Fig. 3C). This suggests the existence of a third, although minor, branch for activating PPO that leads to blackening of the wound site. A summary of these results is shown in Table 1. Together, these results demonstrate that cuticle blackening after clean injury relies strongly on the presence of Hayan, acting through both PPO1 and PPO2. In the absence of Hayan, cuticle blackening can be partially restored by septic injury with the Gram-positive *M. luteus*, and this relies on PPO1 and on the presence of Sp7.

**Table 1.** Melanization intensity after clean injury (CI) or septic injury with M. luteus in adult flies. Intensities ranging from +++ = strong to - = none. Of note: melanization is rescued by M. luteus in Hayan flies. This is dependent on the presence of PPO1.

|  | *w*<sup>111</sup><sub>8</sub> | *Hayan*<sup>SK</sup><sub>3</sub> | *Sp7*<sup>SK</sup><sub>6</sub> | *MP1*<sup>SK</sup><sub>6</sub> | *Hayan*<sup>SK3;PP</sup>*O1*<sup>Δ</sup> | *Hayan*<sup>SK3;PP</sup>*O2*<sup>Δ</sup> | *PPO1*<sup>Δ</sup>; *Sp7*<sup>SK</sup><sub>6</sub> | *PPO2*<sup>Δ</sup>; *Sp7*<sup>SK</sup><sub>6</sub> |
| --- | --- | --- | --- | --- | --- | --- | --- | --- |
| CI | +++ | - | ++ | +++ | - | - | ++ | + |
| *M. luteus* | +++ | + | ++ | +++ | - | + | ++ | + |

### Extracellular components of the Toll pathway are required to activate melanization and to survive *S. aureus* infections

Previous studies have suggested a link between the melanization and Toll pathways (Ligoxygakis, 2002; Matskevich et al., 2010), the latter being critical to resist infection to Gram-positive bacteria in *D. melanogaster* (Leulier et al., 2000; Rutschmann et al., 2000). Currently, how the Toll pathway can impact melanization is not well understood. Our finding that the Gram-positive bacteria *M. luteus* activates blackening in the absence of Hayan prompted us to investigate the role of pattern recognition receptors (PRRs) in melanization. First, we monitored the survival of flies lacking upstream or downstream components of the Toll pathway using our melanization-sensitive *S. aureus* assay. We observed that flies lacking GNBP1 or PGRP-SA, two PRRs implicated in the recognition of Gram-positive bacteria, are as susceptible to *S. aureus* infections as *PPO1^Δ^,2^Δ^* and *Sp7* mutant flies (Fig. 4A). In contrast, flies lacking GNBP3, a PRR sensitive to fungal ß-glucans, resist as wild-type to *S. aureus* infection. Importantly, *spätzle (spz^rm7^*) flies lacking intracellular Toll signaling, exhibit wild-type resistance to *S. aureus* infection (Fig. 4B). This indicates that the high susceptibility of flies lacking PGRP-SA and GNBP1 is not linked to the intracellular Toll pathway, which regulates the Toll transcriptional output. We therefore analyzed the role of the four serine proteases upstream of Spz in the Toll pathway, ModSP, Grass, SPE, and Psh, for their role in resistance to *S. aureus*. Only *ModSP*^1^ and *Grass^Herrade^* mutant flies die at similar rates to *PPO1^Δ^,2^Δ^, GNPB1^Osirìs^* and *PGRP-SA^seml^* flies (Fig. 5A). We conclude that a subset of the extracellular components functioning upstream of the Toll ligand, namely PGRPS-SA, GNBP1, ModSP and Grass, regulate a mechanism of resistance to *S. aureus* that is independent of intracellular Toll signaling.

**Fig. 4:**
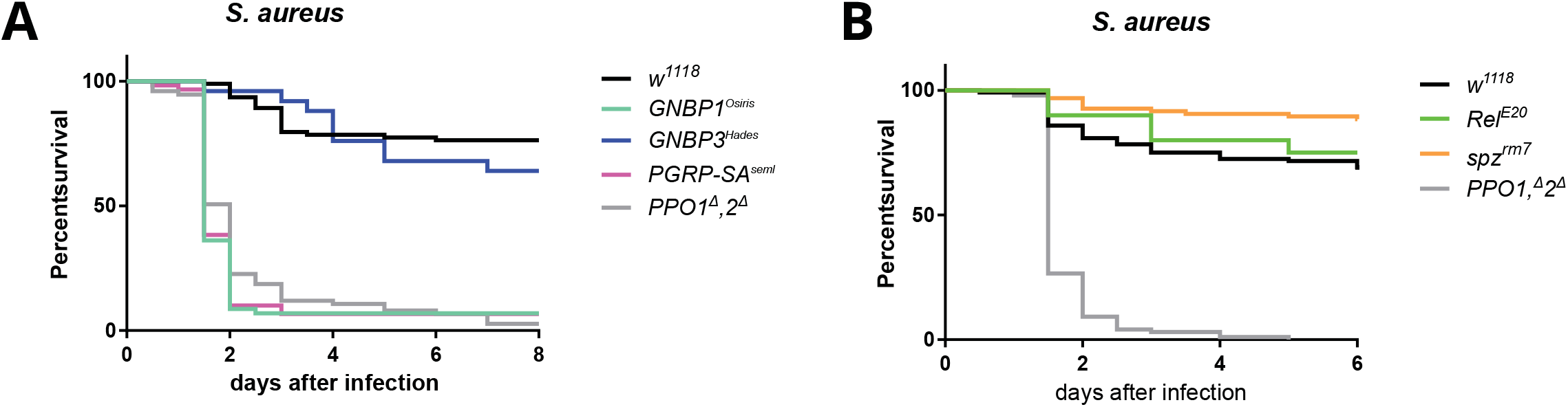
PRRs regulating Toll signaling are critical to resist *S. aureus*. **A:** Survival rates of flies following septic injury with Gram-positive *S. aureus*. Flies lacking *GNBP1* (P < 0.0001) or *PGRP-SA* (P< 0.0001) but not *GNBP3* (P = 0.3141) are less resistant than wild-type flies (w^1118^). **B:** Flies lacking the Toll ligand *Spätzle* (P = 0.0056) or IMD component *Relish* (P= 0.7845) resist *S. aureus* infection similar to wild-type flies (w^1118^). Flies lacking two PPO genes (PPO1^Δ^,2^Δ^) (P < 0.0001) act as the positive control for A and B.

**Fig. 5:**
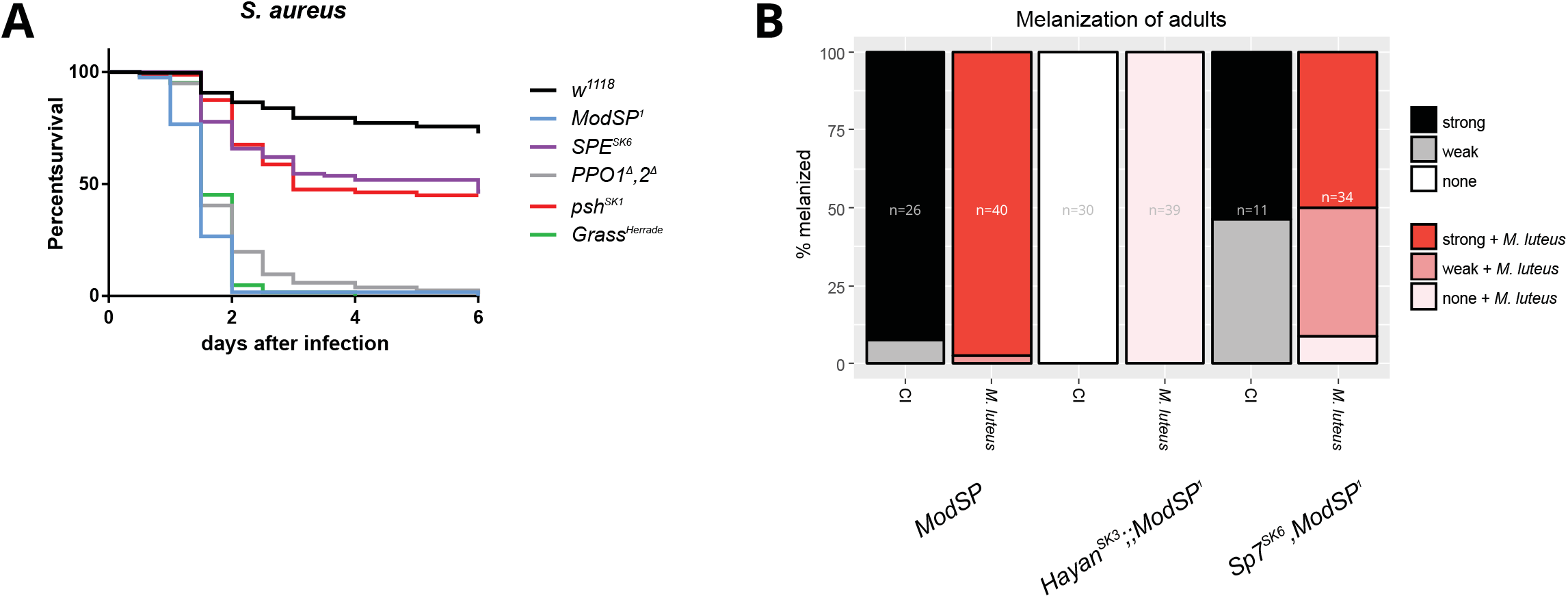
The Toll pathway serine proteases ModSP and Grass are required to survive *S. aureus* infections. **A:** Survival rates of flies following septic injury with Gram-positive *S. aureus*. Flies lacking *ModSP^1^* (P < 0.0001) or *Grass* (P < 0.0001) exhibit high susceptibility compared to wild-type flies (w^1118^). The loss of *psh* (P < 0.0001), SPE or *Spirit* (P < 0.0001) results in a minor survival defect. P-values are from log-rank tests compared with wild-type flies. **B:** More than 90% of *ModSP^1^* flies melanize normally. In contrast, flies mutant for *Hayan^SK3^;;ModSP^1^* do not melanize at all. This effect is dependent on the simultaneous absence of *Hayan* and *ModSP^1^* because *Sp7^SK6^,ModSP^1^* flies do not exhibit a stronger phenotype than *Sp7^SK6^* alone (see Fig. 4). Percentages of total flies are displayed, and n is indicated in each respective bar. Flies were either injured with a clean needle (black bars) or with a needle previously dipped in *M. luteus* solution (red bars). Melanization was assessed in three categories: normal, weak or none (see Fig. S2).

### ModSP/Grass and Hayan contribute independently to different types of melanization

A hypothesis to reconcile our data is that the PRRs GNBP1 and PGRP-SA activate ModSP and Grass, which branch out to activate the melanization pathway. This is consistent with studies in other insects that have shown that melanization is controlled by the same SPs that regulate Toll (An et al., 2009; Kan et al., 2008). Thus, ModSP and Grass could contribute to the blackening reaction, and notably contribute to Hayan-independent cuticle blackening observed upon septic injury by *M. luteus*. To further analyze the role of ModSP in the melanization pathway and its relationship with Hayan and Sp7, we generated *Hayan^SK3^;; ModSP^1^* and *Sp7^SK6^, ModSP^1^* double mutant flies, and compared the blackening reaction at the wound site upon clean and *M. luteus* injury. In agreement with previous studies (Buchon et al., 2009), *ModSP^1^* mutant flies exhibit a wild-type blackening reaction in both conditions. Consistent with our hypothesis that ModSP contributes to Hayan-independent blackening, *Hayan^SK3^;;ModSP^1^* mutants failed to show cuticle blackening regardless of treatment (Fig. 5B). On the contrary, *Sp7^SK6^, ModSP^1^* mutant flies showed a similar level of blackening reaction found in *Sp7* mutants alone (Fig. 5B).

We conclude that at least two independent pathways contribute to cuticle blackening in *D. melanogaster*adults: one involving Hayan and another involving PGRP-SA, GNBP1, ModSP, Grass and Sp7.

### Neither Hayan nor Sp7 alone is required to activate Toll signaling

We found that components of the Toll pathway directly regulate the melanization response. We then tested if the two SPs involved in the melanization pathway, Sp7 or Hayan, could be involved in the activation of the Toll pathway by Spätzle. Consistent with previous studies (Nam et al., 2012; Tang et al., 2006), neither *Sp7* nor *Hayan* single mutants reduce Toll pathway activation after *M. luteus* infection as monitored by the expression of *Drs* (Fig. 6A). Of note, we observed a mild overactivation of the Toll pathway in *Sp7* mutant flies upon *M. luteus* infection compared to wild-type (Fig. 6A). Additionally, no effect on the activation of the Toll pathway could be observed in in *Sp7* nor *Hayan* mutant flies upon *Candida albicans* infection (Fig. S4A). We conclude that Hayan and Sp7 alone are not required for Toll pathway activation by bacteria or fungi.

**Fig. 6:**
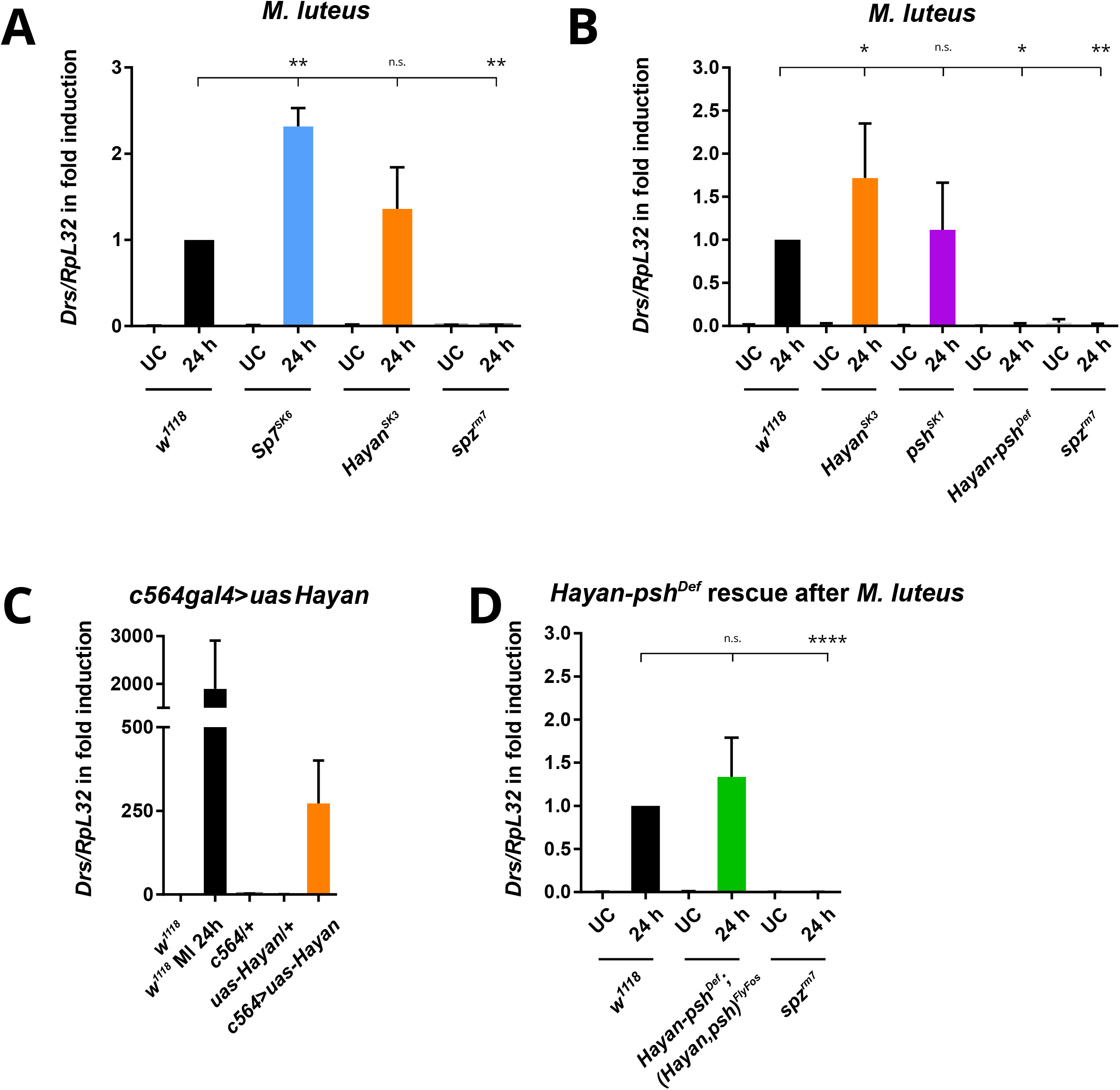
*Hayan* contributes to Toll pathway activation. **A:** Single mutants for *Sp7* and *Hayan* show no loss of Toll pathway activity (*i.e*. expression of *Drosomycin* after *M. luteus* infection). *MP1^SK6^* and *Sp7^SK6^* flies show significantly higher levels of *Drs*, while *Hayan* mutants show no difference after 24 h compared to *w^1118^* flies. **B:** Flies lacking *Hayan* and *psh* simultaneously fail to activate *Drs* expression similar to *spz^rm7^* flies, while single mutants for *Hayan* or *psh* show no significant difference compared to *w^1118^* flies. **C:** Expression of a constitutively active form of *Hayan* in the fat body is sufficient to induce *Drs* in otherwise unchallenged flies. **D:** Insertion of a *Hayan, psh* transgene (FlyFos) rescues to wild-type levels of *Drs* expression in the *Hayan-psh^Def^* double mutant background after *M. luteus* infection. Shown are the relative expression levels of *Drs* relative to *RpL32. w^1118^* levels after 24h (A,B,D) or UC (C) are set to 1. *spz^rm7^* flies act as a negative control. Data analyzed with Mann-Whitney test (A,B, C) or 2 way Anova (E).

### Evidence of a close evolutionary relationship between Hayan and Psh

*Hayan* and the Toll-regulating SP *psh* are only 751 bp apart on the *D. melanogaster* X chromosome. This stimulated us to investigate the possible relationship between Hayan and Psh. We performed a phylogenetic analysis of all *D. melanogaster* CLIP-domain SPs listed in Veillard et al. (2015) using protein sequences from the conserved catalytic domain. We found that Hayan and psh form a closely-related monophyletic lineage amongst CLIP-domain SPs (Fig. 7A). Interestingly, it appears that *psh* is a lineage-restricted duplication found only in Melanogaster group flies, arising from an ancestral *Hayan* gene, the latter being conserved across the genus *Drosophila* (Fig. 7B and Fig. S5A). We found that the Psh protease “bait region”, a region prone to cleavage by pathogen proteases (Issa et al., 2018), is also present in Hayan (Fig. S5B). Furthermore, we recovered a striking similarity in transcript structure of *psh*-RA and *Hayan-*RA, where alternative splicing excludes the fourth exon found in *Hayan*-RB and *Hayan-RD* (Fig. 7C). This 4^th^ exon encodes a “Hayan-exclusive domain,” not found in other *D. melanogaster* CLIP-domain SPs. Nevertheless, this alternative splicing is conserved in FlyBase v2018-02 annotations for *Drosophila pseudoobscura* (subgenus Sophophora) (Fig. S6A), and we independently confirmed this pattern across the genus *Drosophila* using RT-PCR (Fig. S6B). Thus, alternative splicing of *Hayan* is an evolutionarily conserved mechanism to produce *Hayan* transcripts either similar to *psh*, or exclusive to *Hayan*. To shed light on the functional regions of this Hayan-exclusive domain, we extracted *Hayan* sequences from diverse *Drosophila* and used FEL and SLAC to infer codons under selection (Delport et al., 2010). We found multiple sites under purifying selection (p < .05, Fig. S6C, Supplementary Data File 1), forming largely-conserved motifs corresponding to *D. melanogaster Hayan-RB* residues from A^201^-p^206^, r^226^-p^234^, l^261^-v^282^, and D^303^-G^312^. We did not find homologues for the Hayan-exclusive domain outside *Drosophila* in GenBank, UniProt, and Pfam databases. Globally, we conclude that *psh* is a recent duplication of *Hayan*, lacking the Hayan-exclusive domain. Furthermore, transcripts from both genes can encode highly similar proteins.

**Fig. 7:**
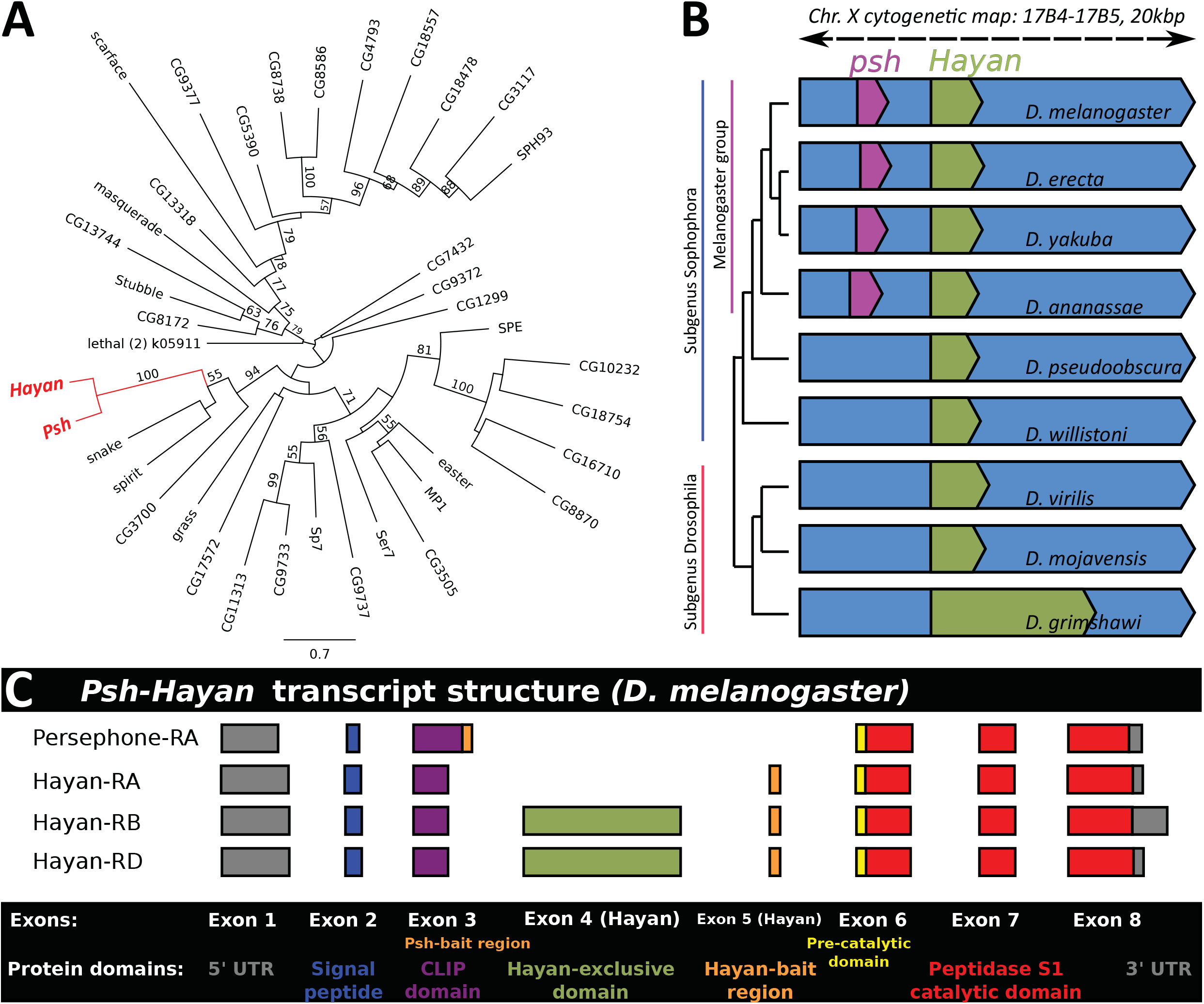
Evolutionarily conserved Hayan isoforms are either similar to Psh, or unique to Hayan. **A:** Maximum likelihood phylogeny of catalytic domains from *Drosophila* CLIP-domain SPs from Veillard et al. (2015). Support values represent 100 bootstraps. Hayan and Psh form a monophyletic lineage within *D. melanogaster* CLIP-domain SPs. **B:** *psh* is a gene duplication restricted to Melanogaster group flies, derived from ancestral *Hayan* (also see Fig. S5 A). Annotations represent CDS gene region*S. Drosophila grimshawi’s Hayan* region is elongated due to a 4000 bp intron between Dgri\GH12343-RB exons 3 and 4. **C:** Hayan-RA and psh-RA bear a striking resemblance in transcript structure, while the *Hayan* transcript isoforms *Hayan-RB* and *Hayan-RD* include a Hayan-exclusive domain not found in other CLIP-domain SPs in *D. melanogaster*.

### Hayan and Psh redundantly regulate Toll pathway activation by the PRR and Psh branch

The duplication producing *Hayan* and *psh* is quite recent raising the possibility that both genes share an overlapping function that would be masked in single mutant analyses. This prompted us to analyze the combined effect of *Hayan* and *psh* mutations by generating a deficiency that removes both genes by CRISPR referred to as *Hayan-psh^Def^*. In these experiments, we also use a newly generated null *psh^SK1^* mutant*S. Hayan-psh^Def^* mutant flies are viable and show no overt morphological defects. Strikingly, we found that *Hayan-psh^Def^* flies fail to activate the Toll pathway upon *M*. *luteus* infection in contrast to single mutants (Fig. 6B). Similarly, *Hayan-psh^Def^* mutants also fail to activate Toll after *C. albicans* infection and *Bacillus sp*. protease injection (Fig. S4B,C). Interestingly, while *Hayan^SK3^* mutants show wild type levels of *Drs* expression after protease injection, *psh^SK1^* flies suppress *Drs* to ~35% of wild-type levels. Only in the simultaneous absence of both Hayan and Psh is *Drs* expression suppressed to *spz^rm7^* levels (Fig. S4C). This suggests that Hayan also contributes to the Psh pathway. Thus, both Hayan and Psh, redundantly regulate the Toll pathway downstream of PRRs. We note that Hayan and Psh cluster phylogenetically with the SP Snake (Fig. 7A), a protease that in dorso-ventral Toll signaling cleaves the terminal SP that cleaves Spätzle (Dissing, 2001; LeMosy et al., 2001; Lindsay and Wasserman, 2014). This suggests Hayan and Psh could play a similar role upstream of SPE. Consistent with these observations, expression of an activated form of *Hayan* using a fat body Gal4 driver (*c564*) leads to upregulation of the Toll readout *Drs* in the absence of infection (Fig. 6C), as previously shown for expression of *psh* (Jang et al., 2006). Finally, re-introduction of the *Hayan-psh* genomic region in double mutants using a Flyfos transgene (Sarov et al., 2016) rescues *Drs* expression to wild-type levels after exposure to *M. luteus* (Fig. 6D). Collectively, our double mutant analysis reveals an unexpected role of Hayan and Psh in the pattern recognition pathway regulating Toll activity.

## Discussion

The melanization reaction in insects is arguably one of the most striking immune reactions, resulting in a visible black spot at the site of infection. While the *Drosophila* immune response has been studied intensely, our knowledge of the melanization reaction has lagged behind. This is in part because *Drosophila* immunity has traditionally focused on immune modules conserved in mammals, but the overwhelming complexity of serine protease cascades has also prevented a clear understanding. Indeed, insect genomes harbor an incredible diversity of serine proteases, whose function remains largely uncharacterized (Cao et al., 2015; Ross et al., 2003). In this study, we provide new insights on the PO cascade and uncover a new relationship between the Toll and melanization pathways.

### Melanization is more than the blackening of the wound area

While the importance of the melanization response has been demonstrated in other insects (*e.g. Manduca sexta* (Eleftherianos et al., 2007; Lu et al., 2008)), the precise relevance of PPOs in *D. melanogaster* host defense was disputed until recently (Ayres and Schneider, 2008; Leclerc et al., 2006; Tang et al., 2006). The use of null mutations in *Drosophila PPO* genes clearly demonstrated that melanization-deficient flies lack resistance against microbes, mainly Gram-positive bacteria and fungi (Binggeli et al., 2014). In this study, we developed an infection model using a low-dose inoculation of the Gram-positive bacteria *S. aureus*, which we find is especially appropriate to study melanization. Strikingly, flies deficient for PPOs, but not flies with impaired AMP production or lacking hemocytes, rapidly succumb to this challenge. The melanization response restricts the growth of *S. aureus* preventing its systemic dissemination. Furthermore, our study shows a disconnect between resistance to infection and the blackening of the wound site. While a mutation in *Hayan* leads to the almost complete loss of the blackening reaction in adults, *Hayan* mutants do not share the susceptibility of *PPO1^Δ^,2^Δ^* flies against *S. aureus*. In contrast, *Sp7* mutant flies do not survive *S. aureus* infection, despite almost wild-type levels of cuticle and hemolymph blackening in adults. This indicates that it is not the blackening *per se* that is involved in the control of *S. aureus*, but rather other reactions downstream of PPO activity. For instance, the melanization response is associated with the production of cytotoxic molecules like ROS (reviewed in Nappi et al., 2009). Consistent with a possible role of ROS, we observed that *Sp7* mutants that are susceptible to *S. aureus* have reduced H_2_O_2_ levels in total fly lysates, but this difference was not significant (Fig. S1E, p = 0.1). It is tempting to speculate that ROS or other cytotoxic intermediates contribute to host resistance to microbial infection, while melanin deposition is involved in host protection by acting as a ROS sink as proposed by other authors (Nappi et al., 2009; Riley, 1997).

### The Toll pathway branches at the level of or downstream of *Grass* to regulate the melanization pathway

Studies in other insects such as *Manduca sexta* and *Tenebrio molitor* have shown that melanization and the Toll pathway share a common upstream activation mechanism that then separates downstream of SPE-like SPs that are capable of cleaving both Spz and PPO (An et al., 2009; Kan et al., 2008; Kim et al., 2008). Prior to our results, there was only indirect evidence in *D. melanogaster* that the Toll pathway activates melanization (Matskevich et al., 2010). It was however noted that Toll regulates many SPs and serpins involved in the melanization cascade at transcriptional levels (De Gregorio et al., 2001; Ligoxygakis, 2002). In addition, over-expression of several SPs functioning upstream of Spz, or a gain-of-function activation of the Toll pathway leads to spontaneous melanization (An et al., 2013; Gerttula et al., 1988; Lemaitre et al., 1995; Tang et al., 2006). Some studies have also reported that Toll PRRs can activate a host defense reaction independent of Toll intracellular signaling, possibly through melanization (Bischoff et al., 2004; Matskevich et al., 2010). Despite these sporadic observations, if and how the PRR-Toll pathway branches to PPO activation remained unknown. Here we show that PGRP-SA, GNBP1, ModSP and Grass, but not SPE, regulate the melanization response after exposure to Gram-positive bacteria. Our double mutant analysis suggests that the PRR cascade diverges downstream of Grass to activate Sp7 and PPO1. Thus, our study demonstrates a direct connection between the extracellular SPs regulating the activation of Toll and the melanization response as observed in other insects. Surprisingly, we found no clear role for MP1 in melanization nor Toll pathway activity, which contradicts a previous RNAi screen that positioned MP1 downstream of Sp7 in the PO cascade (Tang et al., 2006). At this stage, it cannot be excluded that MP1 contributes to melanization in a redundant way. Alternatively, results obtained with RNAi may be the consequence of off-target effects that shut down other SPs.

### Hayan and Psh redundantly regulate Toll signaling downstream of the PRR pathway

The evidence that SPE in other insect species can lead to the activation of both the Toll and melanization pathways led us to verify the roles of two melanization SPs, Hayan and Sp7, in Toll activation. Single mutants for *Hayan* and *Sp7* showed no defect in Toll pathway activation as revealed by the level of *Drosomycin* induction. Surprisingly, we found that Hayan-psh^Def^ mutants fail to activate the Toll pathway regardless of the type of challenge. This points to a crucial function for these two SPs in the activation of the Toll pathway in both the protease and PRR pathways. The Psh bait region motif ^107^GRVDVPTFGS^116^ is critical for protease-dependent Psh cleavage in *Drosophila* S2 cells (Issa et al., 2018). This region is truncated but present in Hayan (Fig. S5), providing a mechanism for microbial proteases to cleave Hayan or Psh differently, leading to microbe-specific activation of downstream Toll. This contraction of the bait region may represent the early beginnings of sub-functionalization amongst *Hayan* and *psh* (He et al., 2005). Indeed, using RT-PCR, the shorter psh-like transcripts from *Hayan* appear less abundant in *D. melanogaster* than outgroup flies where *Hayan* alone should be responsible for Toll signaling (Figure S6C).

We also show that *Hayan* and *psh* arose from a recent duplication of an ancestral *Hayan* gene. Gene duplications can have varied evolutionary outcomes, but the strong reduction of *Drs* expression in *Hayan-psh^Def^* mutants upon septic injury with *M. luteus*, but not in single mutants, suggests Hayan and Psh are redundant in regulating Toll signaling. Additionally, Hayan and Psh share striking similarities in a regulatory region upstream of the catalytic domain, which strongly suggests that they can be cleaved by the same upstream SPs. As both SPs also share strong similarities in their catalytic domain, they likely cleave similar downstream targets. During embryonic dorso-ventral signaling, the SP Snake regulates the Spätzle processing SP Easter (Lindsay and Wasserman, 2014). The phylogenetic relatedness of Snake and Hayan/psh and the relatedness of Easter and SPE (Fig 7A) suggests Hayan and Psh act upstream of SPE. We note that *spz^rm7^* deficient flies have lower *Drs* levels than any other mutants, except *Hayan-psh^Def^* mutants. As *SPE* mutants still express a residual level of *Drs* upon septic injury with *M. luteus* (Fig. S4D), there is likely another SP with the ability to process Spz, as first proposed by Yamamoto-Hino and Goto (2016). CLIP-domain SPs related to Easter and SPE are promising candidates to fulfill this role. Intriguingly, MP1 clusters phylogenetically with Easter, and has been shown to cleave Spz in *Drosophila* S2 cells (Yamamoto-Hino and Goto, 2016). Further, biochemical characterization will be needed to fully clarify the positions of Hayan and Psh in the proteolytic cascade downstream of PRRs.

### Melanization and Toll: more than black and white

Altogether, we propose a revised model of the extracellular SP cascades regulating melanization and the Toll pathway in *D. melanogaster* that takes the three main findings of this work into consideration: i) the existence of two different pathways activating melanization, ii) the involvement of the extracellular PRRs in the melanization reaction and, iii) the implication of both Hayan and Psh in the extracellular PRR and the Psh pathway regulating Toll activity (Fig. 8). According to our model, cuticle injury activates Hayan by an unknown pathway, which results in the deposition of melanin (blackening reaction) around the wound area through both PPO1 and PPO2. After an infection with Gram-positive bacteria, peptidoglycan can be recognized by the PRRs PGRP-SA/GNBP1, leading to an SP cascade involving ModSP, Grass, Hayan and Psh, and SPE, resulting in Spätzle cleavage. In our model, the PRR pathway upstream of Toll branches at the level or downstream of Grass to Sp7, activating PPO1 to combat invading bacteria. Alternatively, microbial proteases and endogenous elicitors (Issa et al., 2018) can activate the Toll pathway independently of PRRs by cleaving Psh, and possibly Hayan, directly. Thus, two SPs, Hayan and Psh merge signals from both the PRR and Psh pathway, to activate a common extracellular pathway upstream of Toll. Globally, we describe a crucial role for Hayan in Toll activation that is performed redundantly alongside Psh, and describe how these proteins act downstream of both PRRs and microbial proteases to activate Toll.

**Fig. 8:**
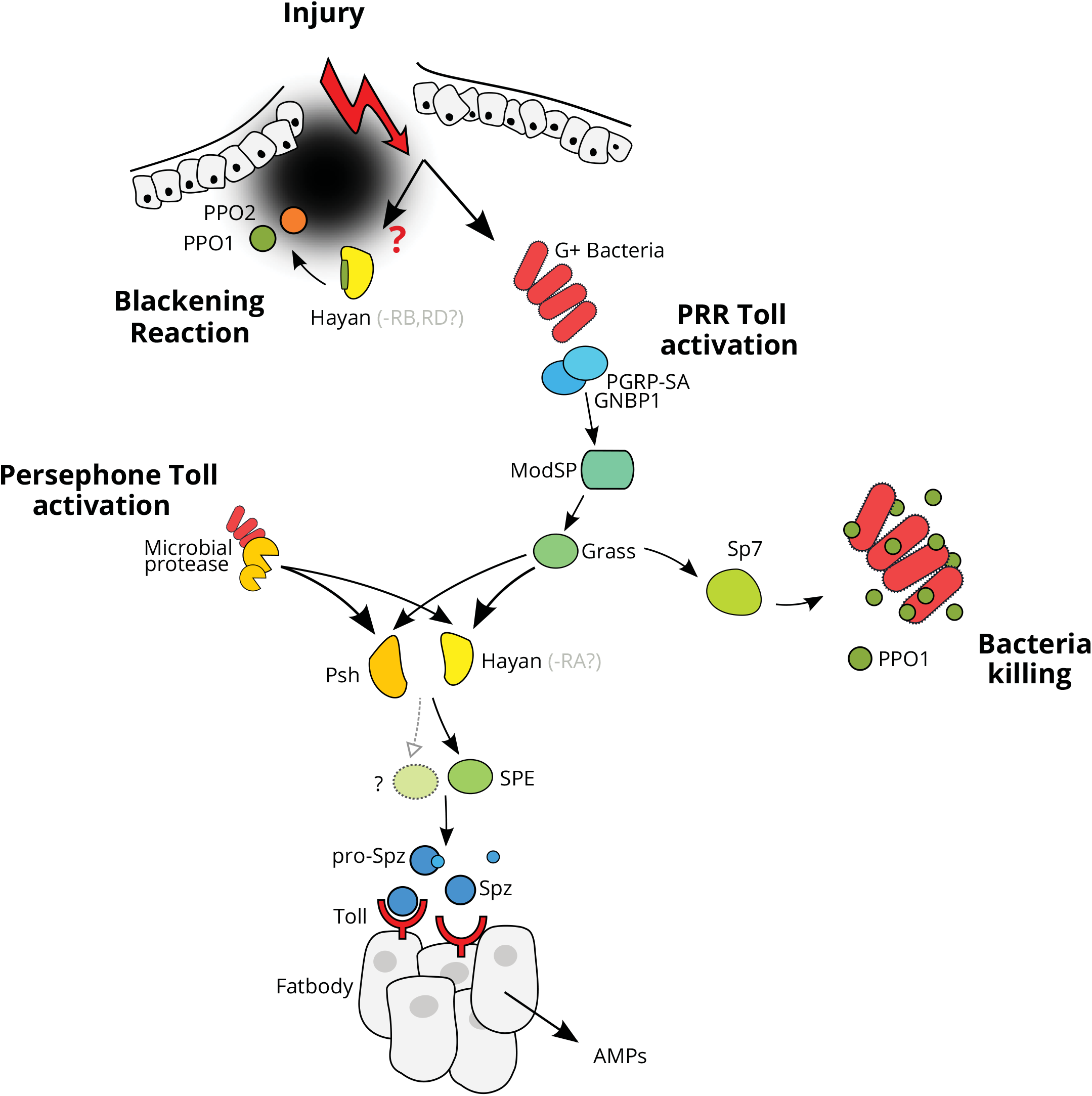
A revised model of SPs regulating the Toll pathway and the melanization reaction. After the introduction of a wound through the cuticle and the underlying epithelium, Hayan can be activated by an unknown mechanism that results in the deposition of melanin around the wound area (left part). This Hayan-dependent blackening reaction can be achieved through both PPO1 and PPO2. If Gram-positive bacteria enter through the wound, peptidoglycan can be recognized by the PRRs PGRP-SA and GNBP1, initiating the sequential activation of the SPs ModSP, Grass Psh/Hayan and SPE. This leads to the cleavage of Spz and the activation of Toll signaling in the in the fat body (middle part). This extracellular SP pathway branches at the level of Grass to Sp7, activating PPO1 to combat invading bacteria, possibly via the production of cytotoxic intermediates but not melanin (right part). Microbial proteases can activate the Toll pathway through the Psh-SPE-Spz extracellular pathway. Although suggested by sequence homology, it is unclear whether microbial proteases can also activate Hayan but both Hayan and Psh regulate the Toll pathway downstream of Grass, ModSP and PRRs. A previous study (Yamamoto-Hino and Goto, 2016) and our data suggest the existence of another SP capable of cleaving Spz beyond SPE.

The analysis of proteolytic cascades regulating the *Drosophila* immune response has been hampered by the large number of SP genes, often found in clusters in the genome. While the biochemical approaches carried out in large insects have allowed a comprehensive understanding of SP signaling cascades in moths and beetles (Kanost and Jiang, 2015), genetic approaches using single gene mutant analysis were unable to determine these cascades in *Drosophila* (Binggeli, 2013). Functional redundancy, as exemplified in our study of *Hayan* and *psh*, clarifies the shortcomings of single-gene genetic approaches. Coupling phylogenetic analysis approaches with double (or triple) compound mutant analysis can pave the way to better characterize these cascades.

## Methods

### Insects Stocks

Unless indicated otherwise, *w^1118^* flies were used as wild-type controls. The *PPO1^Δ^, PPO2^Δ^* and *PPO1^Δ^,2^Δ^, Relish^E20^ (Rel^E20^), spätzle^rm7^ (spz^rm7^), ModSP^1^, Grass^Herrade^* (BL: 67099), *Hayan^SK3^, SPE^SK6^*, *Sp7^SK6^, eater^1^, hml^Δ^-Gal4, c564-Gal4* (BL: 6982), *UAS-bax, UAS-Hayan, GNBP1^osiris^, GNBP3^Hades^, PGRP-SA^seml^* (BL: 55761) lines are described previously or obtained from Bloomington Drosophila Stock Center (Binggeli et al., 2014; Bretscher et al., 2015; Brown et al., 2001; Buchon et al., 2009; Dudzic et al., 2015; Gobert, 2003; Gottar et al., 2006; Nam et al., 2012; Neyen et al., 2014; Sinenko and Mathey-Prevot, 2004; Yamamoto-Hino and Goto, 2016). The *MP1^SK6^, psh^SK1^* and Hayan-psh^Def^ mutant lines were generated by CRISPR/Cas9 as described in (Kondo and Ueda, 2013). *MP1^SK6^* harbors a 13 bp deletion from position 4308655 to 4308668 on the third chromosome. *psh^SK1^* harbors a 10 bp deletion from position 18485863 to 18485873 on the X chromosome. The *Df(1)Hayan,psh^SK5^* deficiency has a 6816 bp deletion from position 18480206 to 18487022 on the X chromosome (referred to as *Hayan-psh^Def^* in the text). To rescue the *Hayan-psh^Def^* mutant flies, transgenic flies were produced by injecting embryos with a *Hayan,psh* transgene from FlyFos (Clone CBGtg9060C0781D, Sarov et al., 2016). This transgene contains ~10 kb of genomic DNA from gene *CG15046* to *Hayan). Drosophila* stocks were maintained at 25°C on standard fly medium.

### Microorganism culture and infection experiments

The bacterial strains used and their respective optical density of the pellet (O.D.) at 600 nm were: the DAP-type peptidoglycan-containing Gram-positive bacteria *Bacillus subtilis (B. subtilis*, O.D 5); the Lys-type peptidoglycan-containing Gram-positive bacteria *Micrococcus luteus (M. luteus*, O.D. 200), and *Staphylococcus aureus (S. aureus*, O.D. 0.5). Strains were cultured in Luria Broth (LB) at 29 °C (*M. luteus*) or 37 °C (other species). The yeast *Candida albicans* ATCC 2001 (*C. albicans*, OD 400) was cultured in YPG medium at 37 °C. Pellets were diluted in distilled water. The *S. aureus-* GFP strain is described in Needham (2004). Systemic infections (septic injury) were performed by pricking adults in the thorax with a thin needle previously dipped into a concentrated pellet of bacteria. Infected flies were subsequently maintained at 29 °C (*M. luteus, C. albicans, B. subtilis*) or at 25 °C (*S. aureus*, injection of *B. subtilis* protease). At least three tubes of 20 flies were used for each survival experiment and survival was scored once or twice daily. For lifespan experiments, flies were kept on normal fly medium at 25 °C and were flipped every two days. 18 nL of protease of *Bacillus sp*. (Sigma P0029) diluted 1:1500 in PBS was injected into the thorax for qRT-PCR experiments.

### Wounding experiment

Clean injury (CI) refers to an injury performed with an ethanol sterilized needle. A low level of bacterial contamination is still possible since the surface of the insect was not sterilized. For imaging of the blackening reaction upon pricking, the thorax of the animal was pricked (as described in infection experiments) using a sterile needle (diameter: ~5 μm). Pictures were taken 16 hours post-pricking. Third instar larvae were pricked dorsally near the posterior end, using a sterile needle (diameter: ~5 μm). Pictures of melanized larvae were taken one-hour post-injury. Pictures were captured with a Leica M 205 FA microscope, a Leica DFC7000FT camera and the Leica Application Suite. For publication purposes, brightness and contrast were increased on some images.

### Melanization assessment

Flies or larvae were pricked as described and the level of blackening at the wound site, estimated by the size and color of the melanin spot, was examined 16-18 h later in adults and 3 h later in larvae. For observing the general capacity of hemolymph to melanize, hemolymph was collected from third instar (L3) larvae by dissection and transferred to a 96 well microtiter plate. After incubation at room temperature blackening was recorded by taking pictures.

### Bacterial load of flies

Flies were infected with *S. aureus* as described above. At the appropriate time point, flies were sacrified by washing them in 70% ethanol. This treatment is expected to remove bacteria from the surface of the flies. Ethanol was washed off with sterile PBS and groups of five flies were homogenized with a PRECELLYS™ homogenizer in 0.2 ml PBS. The homogenate was serially diluted and plated on LB agar. After incubation at 37 °C overnight, colonies were counted and calculated to single fly CFUs. Only *S. aureus* bacterial colonies were recovered at this time point with this approach.

### H_2_O_2 assay_

H2O2 levels were assessed with Fluorimetric Hydrogen Peroxide Assay Kit (Sigma MAK165) according to the manufacturer’s documentation. Briefly, seven adult flies were homogenized in 0.12 ml PBS with a PRECELLYS and then centrifuged at 4°C and 13,000 RPM. Afterwards 0.1 ml homogenate was transferred into a new tube. Sample volumes of 30 μl were used for the assay in duplicate. Protein concentration of the samples was determined using the Bradford assay and results were normalized to the respective protein levels.

### Quantitative RT-PCR

For quantification of mRNA, whole flies or larvae were collected at indicated time points. Total RNA was isolated from 10-15 adult flies by TRIzol reagent and dissolved in RNase-free water. 0.5 micrograms total RNA was then reverse-transcribed in 10 μl reactions using PrimeScript RT (TAKARA) with random hexamer and oligo dT primers. Quantitative PCR was performed on a LightCycler 480 (Roche) in 96-well plates using the Applied Biosystems™ SYBR™ Select Master Mix. Primers were as follows: *Diptericin* forward 5′-GCTGCGCAATCGCTTCTACT-3′, reverse 5’-TGGTGGAGTGGGCTTCATG-3’; *Drosomycin* forward 5’-CGTGAGAACCTTTT CCAAT AT GAT-3’, reverse 5’-TCCCAGGACCACCAGCAT-3’; *RpL32 (Rp49*) forward 5’-GACGCTTCAAGGGACAGTATCTG-3’, reverse 5’-AAACGCGGTTCTGCATGAG-3’.

### SP sequence analysis

A list of *Drosophila melanogaster* CLIP-domain serine proteases was generated from Veillard et al. (2015), and all *D. melanogaster* transcript isoforms were extracted from FlyBase v2018_02 (Gramates et al., 2017). Translated catalytic domains for these SPs were aligned using MAFFT (Katoh and Standley, 2013) for maximum likelihood phylogenetic analysis using PhyML in Geneious 10.2.3 (Guindon et al., 2010; Kearse et al., 2012). Using the FlyBase genome browser, 100kb gene regions surrounding Hayan orthologues were extracted from various *Drosophila* species, aligned using Hayan as a frame of reference, and manually searched for conserved SP motifs (*e.g*. “LTAAHC”) common to all *D. melanogaster* CLIP-domain SPs. Following Hayan and Psh characterization, annotated *Hayan* transcripts were extracted from FlyBase and recent immune annotations of subgenus *Drosophila* flies (Hanson et al., 2016). Using these genomic and transcriptomic data, a Hayan-exclusive domain was extracted from diverse *Drosophila* and analyzed for signatures of selection using FEL and SLAC analyses implemented in datamonkey.org (Delport et al., 2010; Kosakovsky Pond and Frost, 2005). We also performed these analyses for comparisons of Hayan to Psh. Supplementary Data File 1 accessible at http://bit.ly/2LKintJ.

### Statistical analysis

Each experiment was repeated independently a minimum of three times (unless otherwise indicated), error bars represent the standard deviation (s.d.) of replicate experiments (unless otherwise indicated). Statistical significance of survival data was calculated with a log-rank test compared to wild-type flies, and P-values are indicated in figure legends. Statistical significance of qPCR or ROS data was calculated with Two-way ANOVA or Mann-Whitney test. Melanization data was analyzed with Pearson’s Chi-squared test. P-values of < 0.05 = *, < 0.005 = **, and < 0.0005 = *** were considered significant.

## Authors’ contributions

JD and BL designed the study. JD performed the experiments. MH performed bioinformatic analyses. JD, MH and BL analyzed the data and wrote the manuscript. II and SK supplied critical reagents. All authors read and approved the final manuscript.

## Acknowledgements

We thank our colleagues Jean-Phillipe Boquete for injecting fly embryos, Claudine Neyen for comments on the manuscript, Dylan Main for technical support and the Bloomington Drosophila Stock Center for fly stocks.

---

**Fig. S1:**
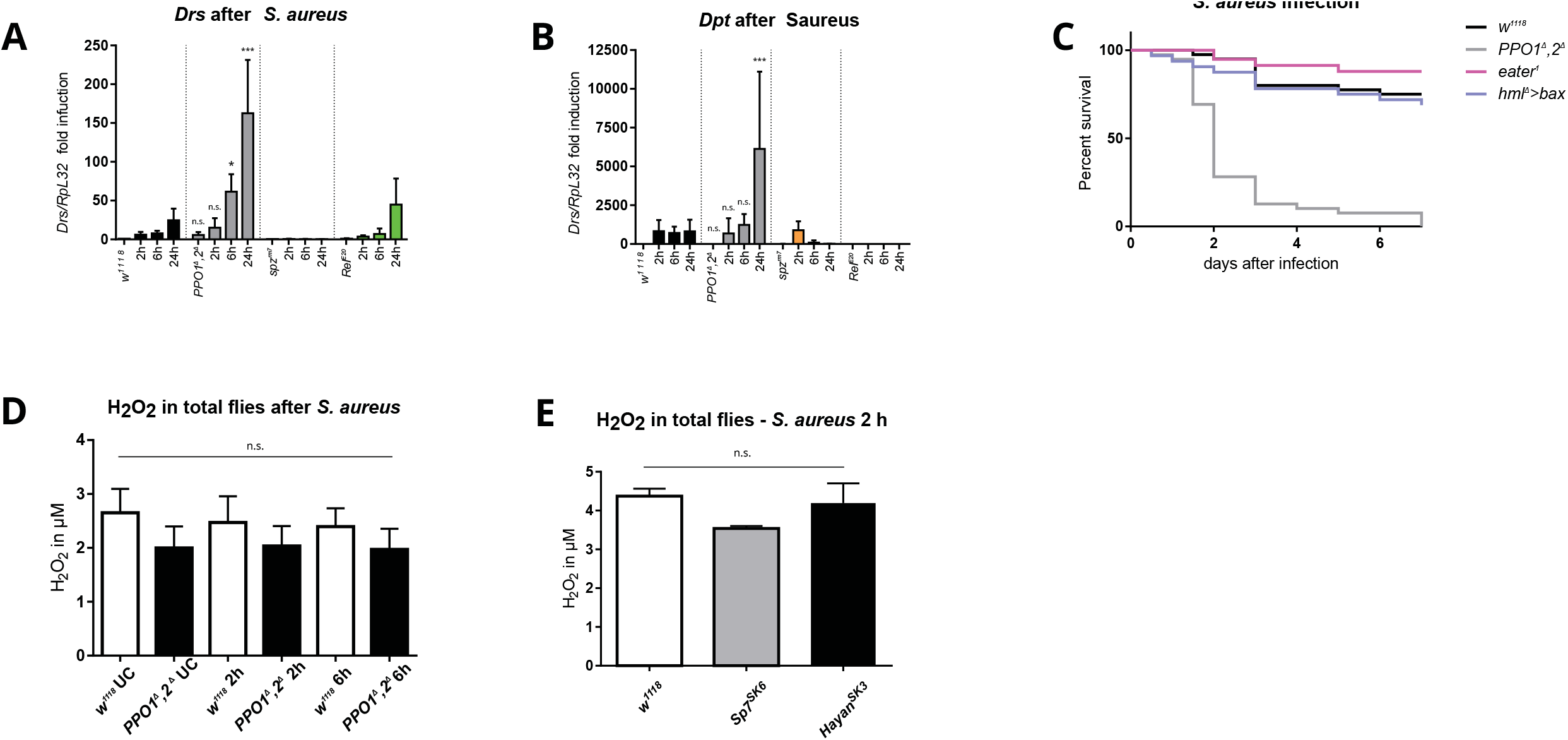
Resistance against low dose *S. aureus* infections is strongly dependent on melanization. **A,B:** *PPO1^Δ^,2^Δ^ flies* show reduction of Toll and IMD pathway activity. Expression of *Drosomycin* (A) or *Diptericin* (B) are given after septic injury with *S. aureus*. Shown are the expression levels of *Drs* or *Dpt* relative to *RpL32. PPO1^Δ^,2^Δ^ flies* show higher levels of *Drs* after 6 and 24 h, and *Dpt* after 24 h compared to *w^1118^* flie*S. Spätzle^rm7^* and *Relish^E20^* flies act as negative control. Data were analyzed with two-way ANOVA, only P-values for *PPO1^Δ^,2^Δ^* compared to respective *w^1118^* timepoints are indicated. Values represent the mean ± standard deviation of at least three independent experiments. **C:** Survival rates of flies following septic injury with Gram-positive *S. aureus*. Flies lacking the hemocyte receptor *Eater*, important for phagocytosis of *S. aureus* (P = 0.10) or flies lacking plasmatocytes by overexpressing the pro-apoptic gene *bax* (P = 0.5118) are as resistant as wild-type flies (w^1118^). P-values are from log-rank tests compared to wild-type flies. **D:** H2O2 measured in whole fly lysates from flies infected with *S. aureus* after 2 and 6 hours compared to unchallenged flies. No significant differences were found between *PPO1^Δ^,2^Δ^* and *w^1118^* samples, although *PPO1^Δ^,2^Δ^* samples show a trend towards lower values. **E:** H_2_O_2_ measured in whole fly lysates from flies infected with *S. aureus* after 2 hours. No significant differences were found between *w^1118^, Sp7^SK6^* and *Hayan^SK3^* samples, although *Sp7^SK6^* samples have lower values. Data analyzed with two-way ANOVA for D and E.

**Fig. S2:**
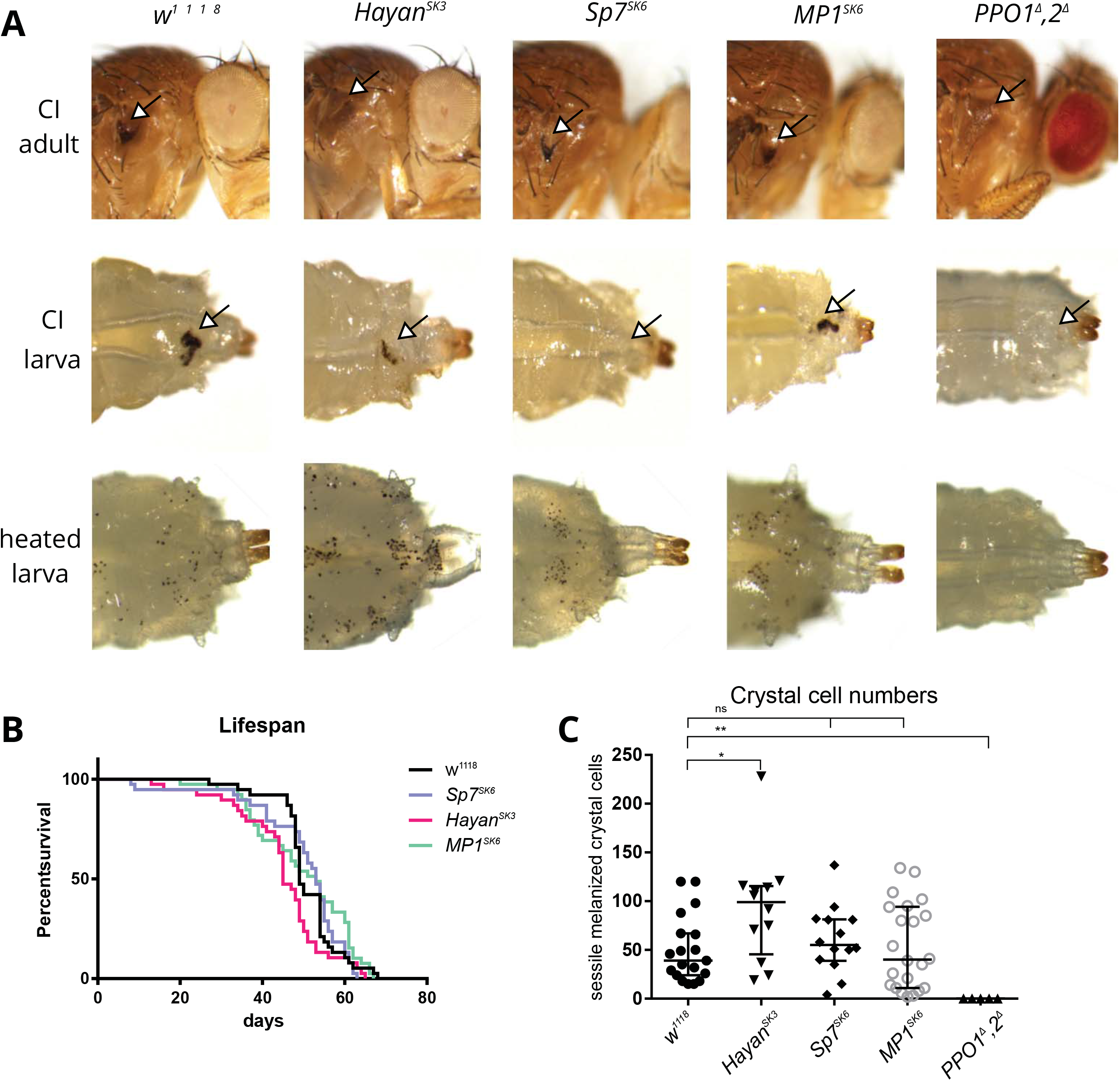
*MP1* deficient flies show no over defect in Melanization. **A:** Blackening of the wound area after clean injury in adult flies is only abolished in *Hayan^SK3^* and *PPO1^Δ^,2^Δ^* flies, while in larvae *Hayan* mutants show a strong defect and *Sp7* mutants a total loss of melanization. All SP mutants show melanized crystal cells after heating. **B:** No severe lifespan defect can be observed in all SPs mutant flies compared to *w^1118^. Hayan^SK3^* flies show a slight reduction in the median survival value from *w^1118^* 49 days to *Hayan^SK3^* 45 days (P = 0.0186), while differences between *MP1^SK6^* or *Sp7^SK6^* and *w^1118^* do not reach significance. The x-axis is the survival time in days and the y-axis is the percentage of living flies. P-values from log-rank test. **C:** Crystal cell counts after heating L3 larvae reveal no significant differences between *Sp7^SK6^* or *MP1^SK6^* and *w^1118^* numbers, while *Hayan^SK3^* crystal cell numbers are around twice as high compared to wild type. Data analyzed with unpaired t-test.

**Fig. S3:**
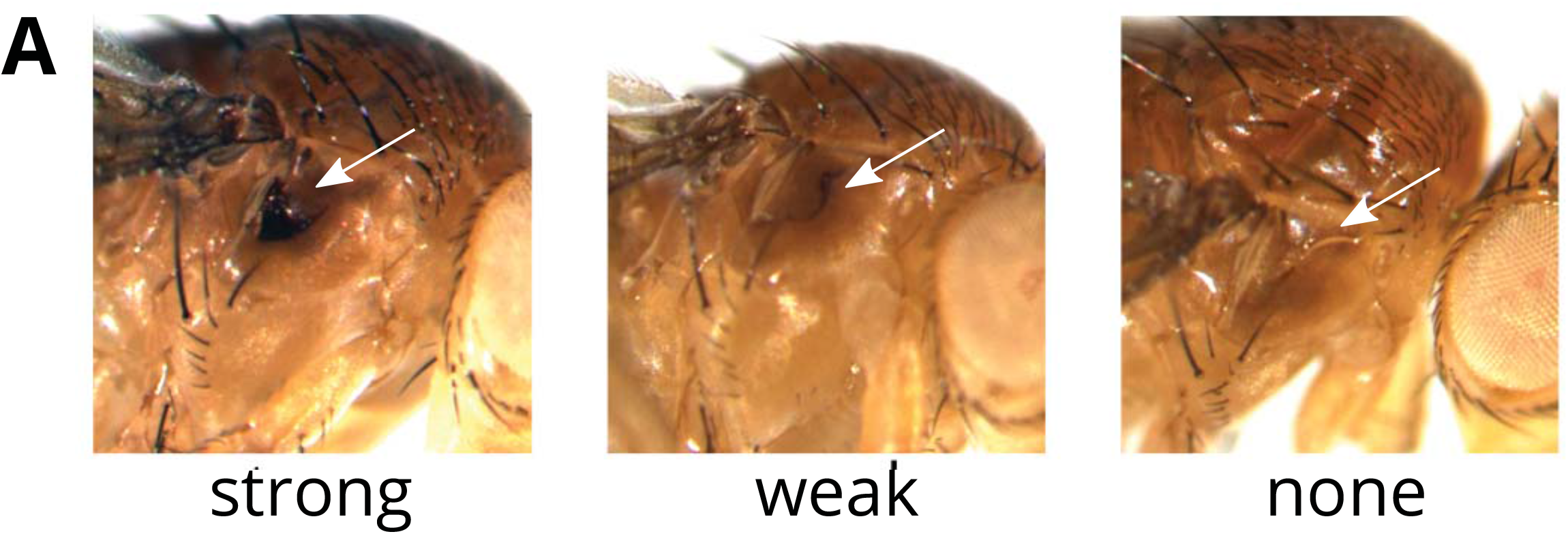
Illustration of melanization intensities used for categories. Flies (*w^1118^:* strong, weak or *PPO1^Δ^,2^Δ^:* none) were injured with a clean needle. Intensities are categorized as normal, weak, or no melanization.

**Fig. S4:**
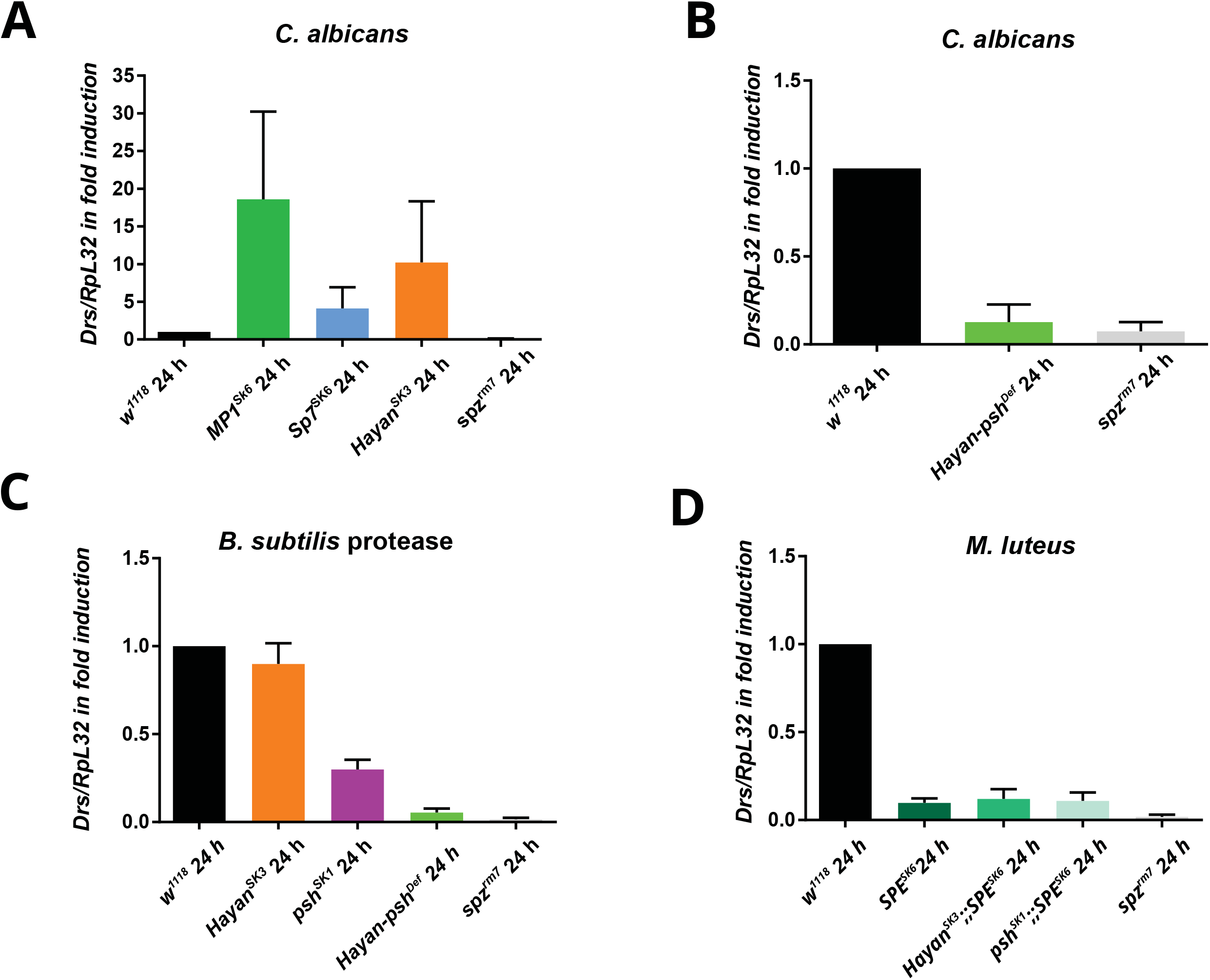
*Hayan, psh* mutant flies have reduced *Drs* expression upon *C. albicans* or protease injection. **A:** Neither *MP1^SK6^, Sp7^SK6^* nor *Hayan^SK3^* fail to activate *Drs* expression after being infected with *C. albicans* relative to *w^1118^* induction (set to 1). **B:** *Hayan,psh* mutants fail to activate the Toll pathway after *C. albicans* infection. **C:** *Hayan^SK3^* shows *w^1118^* levels of *Drs* expression after injection of purified *Bacillus sp*. protease. In contrast, *psh^SK1^* flies suppress *Drs* to ~35% of wild-type level*S. Drs* expression in *Hayan-psh^Def^* is suppressed to a level similar to *spz^rm7^* flies. **D:** *SPE^SK6^, Hayan^SK3^;;SPE^SK6^* and *psh^SK1^;;SPE^SK6^* mutants block *Drs* expression but not as completely as *spz^rm7^* flies. Shown are the expression levels of *Drs* relative to *RpL32. spz^rm7^* flies act as a negative control. Results for three (A,B,D) or two (C) individual experiments are shown.

**Fig. S5:**
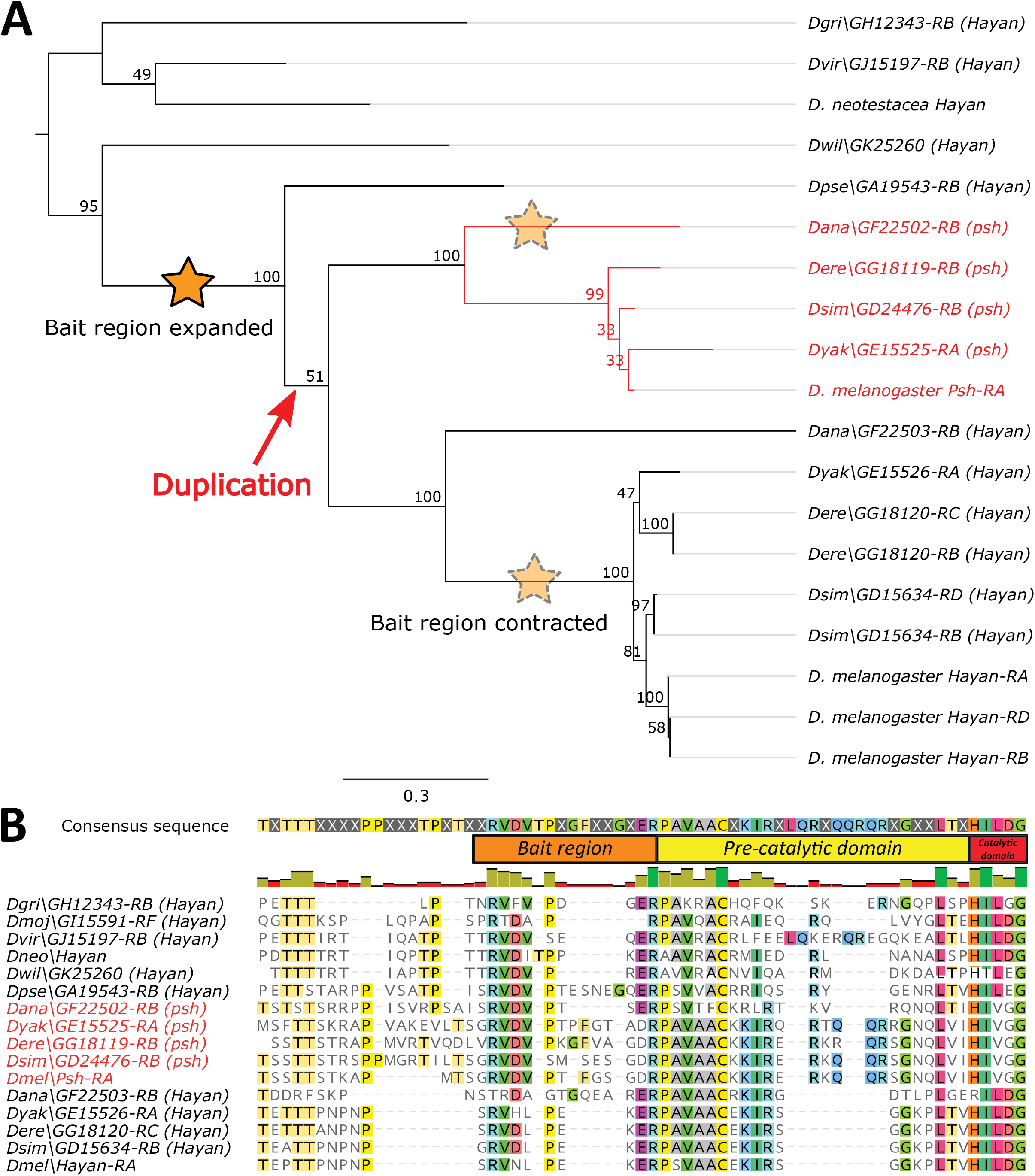
Hayan and Psh are two related serine proteases. **A:** Maximum likelihood phylogeny of unique CDS regions from *Drosophila* Hayan and Psh isoforms. Support values represent 100 bootstraps. Psh forms a paraphyletic lineage nested within *Drosophila* Hayan, with branching patterns largely matching known sorting of *Drosophila* groups. Orange stars indicate the expansion of the microbial protease bait region seen in *D. melanogaster* Psh ^114^FGS^116^ (Fig. S5 B), and translucent stars indicate reversion to the ancestral Hayan bait region similar to *D. melanogaster* Hayan. **B:** Hayan also encodes the protease bait region described by Issa et al. (2018). The important bait motif corresponding to Psh ^108^RVDVP^112^ is largely conserved in outgroup Hayan sequences, while Melanogaster group Hayan has diverged to an RVXLP motif. Interestingly, *D. ananassae* Psh has lost the expanded bait region (following the RVDVP motif), instead retaining these extra cleavage sites in *D. ananassae* Hayan. Following this bait region, the Hayan and Psh pre-catalytic domain is remarkably well-conserved, with some lineage-specific additional sites directly upstream of the Peptidase S1 catalytic domain.

**Fig. S6:**
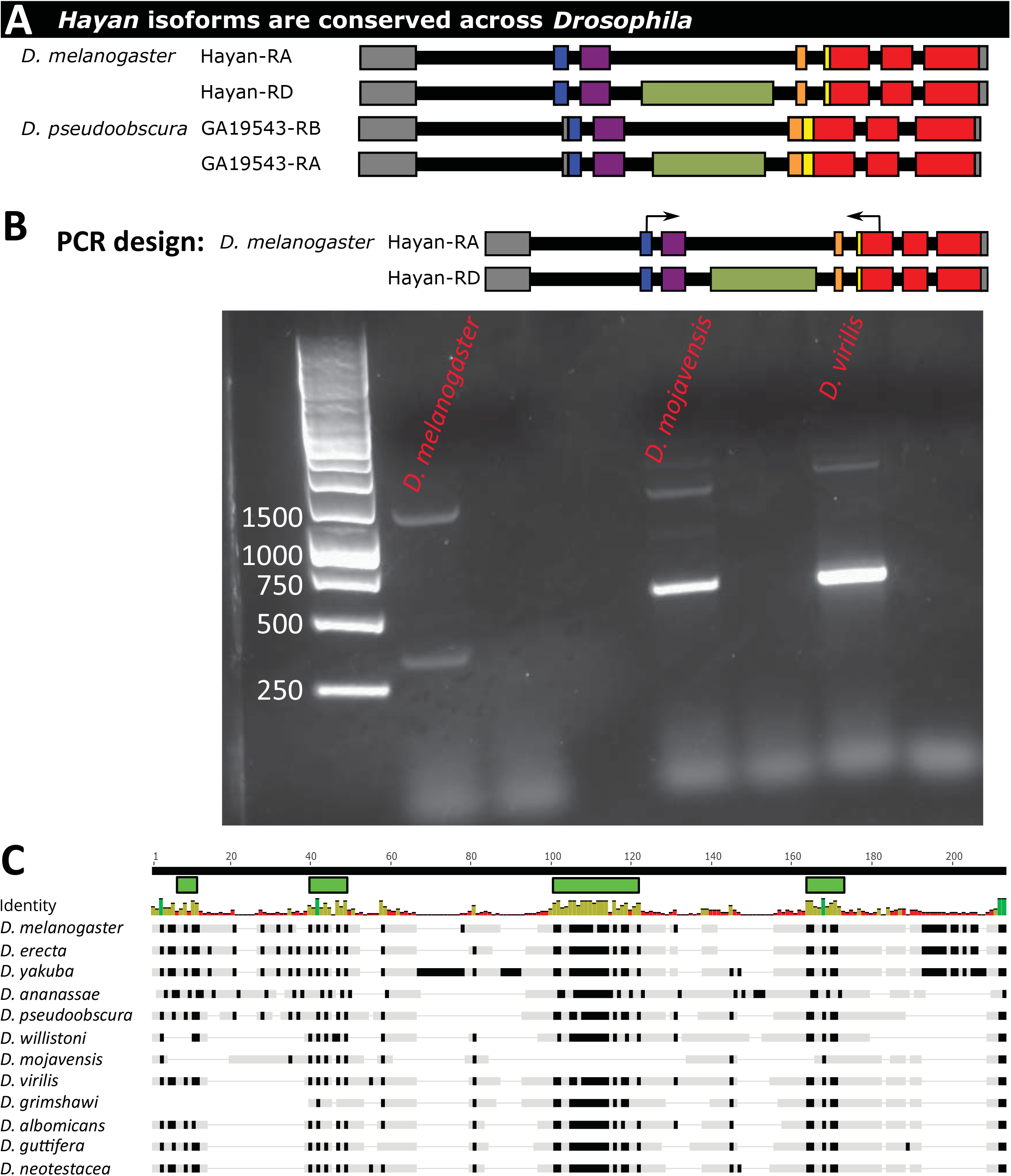
Alternative isoforms of *Hayan* are conserved throughout *Drosophila*. **A:**Alternate *Hayan* isoforms are conserved outside the Melanogaster group in Flybase annotations for *D. pseudoobscura GA19543*. **B:** Independent RT-PCR of *Drosophila* species cDNA shows alternate *Hayan* isoforms in *D. melanogaster* and the subgenus Drosophila flies *D. mojavensis* and *D. virilis*. **C:** Codon-aligment of the Hayan-exclusive domain from diverse *Drosophila*. Green bars represent motifs with many residues under purifying selection (Supplementary Data File 1), which also display strong conservation of sequence (black residues in alignment). Notably, *Drosophila mojavensis* lacks most conserved motifs in its Hayan-exclusive domain, and this region is difficult to interpret with available sequence data in the related *Drosophila buzzatii*.

